# Microbiota-derived TMAO links PM_2.5_ exposure to cognitive impairment: A gut–liver–brain axis

**DOI:** 10.64898/2026.05.24.727565

**Authors:** Akhlaq Hussain, Yunxiao Zhong, Shicong Du, Zonghao Ma, Tong Cheng, Jiasui Yu, Lee Cheuk Hin, Douglas Affonso Formolo, Patrick K. H. Lee, Baomin Wang, Kenneth King-yip Cheng, Siu-Wai Wan, Daniel Kam-wah Mok, Jiachi Chiou, Xiangdong Li, Hai Guo, Suk-yu Yau

**Affiliations:** Department of Rehabilitation Sciences, Faculty of Health and Social Sciences, The Hong Kong Polytechnic University, Hung Hom, Hong Kong S.A.R., China; Research Institute of Smart Ageing (RISA), The Hong Kong Polytechnic University, Hung Hom, Hong Kong S.A.R., China; Research Institute for Future Food (RiFood), The Hong Kong Polytechnic University, Hung Hom, Hong Kong SAR., China; School of Energy and Environment, City University of Hong Kong, Hong Kong., SAR China; State Key Laboratory of Marine Environmental Health, City University of Hong Kong, Hong Kong SAR., China; Climate Impact and Environmental Resilience Research Center, City University of Hong Kong, Hong Kong SAR., China; Department of Health Technology and Informatics, The Hong Kong Polytechnic University, Hong Kong SAR., China; Shenzhen Research Institute, The Hong Kong Polytechnic University, Hong Kong., SAR China; Department of Food Science and Nutrition, Faculty of Science, The Hong Kong Polytechnic University, Hung Hom, Hong Kong SAR., China; Department of Civil and Environmental Engineering, The Hong Kong Polytechnic University, Hung Hom, Kowloon, Hong Kong SAR., China; CAS GIG-PolyU Joint Laboratory of the Guangdong-Hong Kong-Macao Greater Bay Area for the Environment; Hong Kong Branches of Chinese National Engineering Research Centers, Polytechnic University, Hung Hom, Hong Kong SAR., China; Research Institute for Land and Space, The Hong Kong Polytechnic University, Hung Hom, Hong Kong S.A.R., China

**Keywords:** PM_2.5_, Gut dysbiosis, Trimethylamine N-oxide (TMAO), Endoplasmic reticulum stress, Hippocampal impairment

## Abstract

Although fine particulate matter (PM_2.5_) is linked to cognitive impairment, the mechanistic links between pulmonary exposure and neurodegeneration remain poorly understood. This study investigates the role of the gut–liver–brain axis in mediating PM_2.5_-induced neurotoxicity. We demonstrate that three weeks of intratracheal PM_2.5_ instillation in mice causes significant memory deficits, impaired hippocampal adult neurogenesis, and reduced synaptic plasticity in both sexes. Metagenomic analysis revealed that PM_2.5_ alters gut microbiota composition, specifically by upregulating pathways involved in trimethylamine (TMA) synthesis. This microbial shift led to a systemic increase in the metabolite trimethylamine N-oxide (TMAO) and its accumulation in the hippocampus. Mechanistic experiments revealed that TMAO drives neurotoxicity by activating the hippocampal PERK-mediated endoplasmic reticulum (ER) stress pathway. Critically, these deficits were reversed by suppressing hepatic TMAO production via either pharmacological inhibitors or genetic knockdown. Additionally, hippocampal-specific PERK silencing or dietary resveratrol intervention attenuated hippocampal ER stress, thereby protecting against PM_2.5_-induced cognitive and hippocampal impairments. These results identify the microbial metabolite TMAO as a key mediator of air pollution-related neurotoxicity and highlight the gut–liver–brain axis as a promising therapeutic strategy to counteract the neurological impacts of air pollution.

## Introduction

Air pollution is a major global health threat, and fine particulate matter (PM_2.5_) contributes substantially to mortality and disability worldwide ^1^. Although the respiratory and cardiovascular risks of PM_2.5_ exposure are well established ^2–4^. Recent studies also indicate substantial effects on the central nervous system, including increased susceptibility to Alzheimer’s disease (AD), in mice ^5^. Importantly, animal models show that PM_2.5_ impairs hippocampal neurogenesis and cognitive function ^6–8^. However, the mechanisms linking PM_2.5_ exposure to neuronal deficits remain poorly understood.

Recent research suggests that PM_2.5_ can affect the brain both directly, by disrupting the blood-brain barrier (BBB) ^9^, and indirectly ^10^, through peripheral mechanisms such as systemic inflammation and the gut-brain axis (GBA) ^11^. GBA is a bidirectional communication network between the gut microbiota and the brain ^12^, is increasingly recognized as a critical modulator of neurological health through changes in gut microbial-derived metabolites ^13^, some of which can cross the BBB to directly affect neuronal function ^14^. Despite increasing interest in the gut-brain axis in neurological disease, its contribution to PM_2.5_-induced neurotoxicity remains poorly understood ^15,16^. This gap may be addressed through further investigation of gut microbiota-derived metabolites, particularly trimethylamine N-oxide (TMAO). TMAO is produced when gut bacteria metabolize dietary nutrients such as choline, betaine, and L-carnitine into trimethylamine (TMA) in the gut ^17^, which is subsequently oxidized in the liver ^18^.

Beyond its established links to cardiometabolic disease, TMAO has recently been implicated in neurological dysfunction and behavioral abnormalities ^19,20^. Mechanistically, TMAO can activate the protein kinase R-like endoplasmic reticulum kinase (PERK) pathway and promote endoplasmic reticulum (ER) stress ^21,22^, a process closely associated with impaired synaptic function, reduced neurogenesis, and cognitive decline. Because PM_2.5_ exposure is also known to alter gut microbial composition ^23^, it is plausible that PM_2.5_ may increase TMAO production and thereby engage ER stress pathways in the brain.

Here, we tested the hypothesis that chronic PM_2.5_ exposure impairs memory and hippocampal integrity through a gut microbiota–derived TMAO/PERK signaling axis. Using a mouse model of chronic PM_2.5_ exposure, we examined behavioral performance, hippocampal neurogenesis, gut microbial composition and function, circulating TMAO levels, and ER stress signaling. We further evaluated causality by inhibiting TMAO production and by selectively reducing hippocampal PERK expression, and we assessed whether a resveratrol-enriched diet could mitigate these effects. Together, this study defines a gut-brain mechanism linking PM_2.5_ exposure to hippocampal dysfunction and identifies TMAO as a potentially actionable target in environmental neurotoxicity.

## Materials and methods

### Animal maintenance and treatment

Both male and female mice, aged 5 to 6 weeks, were housed under standard laboratory conditions with ad libitum access to chow and water, maintained on a 12-hour light/dark cycle. The mice were randomly assigned to either a control group or a PM_2.5_ exposure group. Mice in the exposure group received intratracheal instillations of 30□μL of PM_2.5_ suspension (2.5□μg/μL in artificial lung fluid) three times weekly for a duration of three weeks, resulting in a dose of 3 mg/kg body weight per administration for a standard 25 g mouse. This dosage is intermediate between the low dose (1 mg/kg) and the high dose (5 mg/kg) employed by Ku et al. (2017) ^24^. The control group received intratracheal instillations of 30 μL of vehicle (artificial lung fluid, ALF) only. All experimental procedures were reviewed and approved by the Animal Subjects Ethics Sub-Committee of The Hong Kong Polytechnic University and conducted in accordance with institutional guidelines. Following the exposure period, mice underwent behavioral assessments to evaluate memory performance, as well as immunohistochemical analyses and electrophysiological recordings to investigate hippocampal neurogenesis and synaptic plasticity (Fig. 1a).

**Fig. 1.**
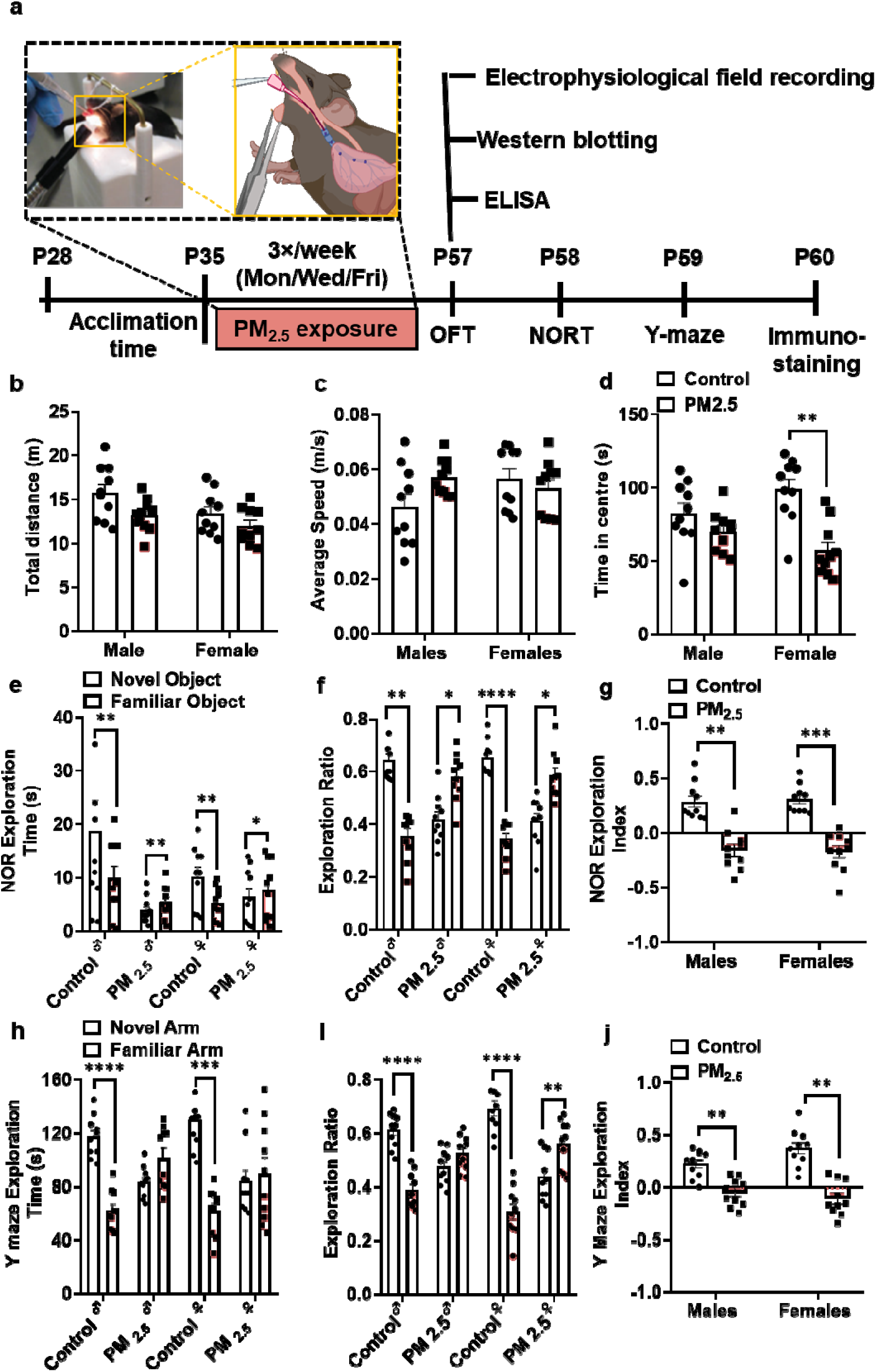
Chronic PM2.5 exposure impairs memory performance. **a** Experimental design. Mice received intratracheal instillation of PM2.5 or artificial lung fluid (ALF) every other day for 3 weeks beginning at postnatal day 35 (P35). **b–d** Open field test (OFT). **b** Total distance traveled (males, P=0.0852; females, P=0.6712). **c** Average speed (males, P=0.0937; females, P=0.8897). **d** Time spent in the center (males, P=0.4490; females, **P=0.0036). **e–g** Novel object recognition test (NORT): Control mice of both sexes showed a significant preference for the novel object, whereas PM_2.5_-exposed mice showed impaired recognition memory. **e** Exploration time (control: males, **P=0.0020; females, **P=0.0018; PM2.5: males, *P=0.0044; females, **P=0.0277). **f** Exploration ratio (control: males, **P=0.0020; females, ****P=0.0001; PM2.5: males, *P=0.0201; females, *P=0.0079; paired t-tests). **g** PM2.5 impaired recognition memory in both sexes. Two-way ANOVA of the exploration index revealed a significant main effect of treatment [F(1, 9) = 103.4, P < 0.0001], with no effect of sex [F(1, 9) = 0.03, P=0.87] and no sex × treatment interaction [F(1, 9) = 0.08, P = 0.79]. Tukey’s multiple-comparisons test confirmed significant reductions in both males (P=0.0002) and females (P=0.0025) following PM2.5 exposure. **h-j** Y-maze test: Control mice exhibited a clear preference for the novel arm, whereas this preference was absent after PM2.5 exposure. **h** Exploration time in the novel and familiar arms. Paired t-test showed that the control males (****P=0.0008) and females (***P=0.00040) spent significantly more time in the novel arm than in the familiar arm. In contrast, PM2.5-exposed male mice (P=0.0667) and PM2.5-exposed female mice (P=0.7300) showed no significant arm preference. **i** Exploration ratio in the Y-maze (control: males, ****P=0.0001; females, ****P=0.0001; PM2.5: males, P = 0.1068; females, **P=0.0059; paired t-tests). **j** PM2.5 impaired spatial working memory independently of sex. Two-way ANOVA of the Y-maze exploration index revealed a significant main effect of treatment [F(1, 9) = 65.01, P=0.0001], with no significant effect of sex [F(1, 9) = 2.07, P=0.18] and no sex × treatment interaction [F(1, 9) = 3.36, P=0.10]. Tukey’s multiple-comparisons test showed significant differences between control and PM2.5 groups in both males (**P=0.0059) and females (**P=0.0014). n=10/group. All data are expressed as mean ± SEM.

### PM_2.5_ sampling and preparation

Roadside ambient PM_2.5_ samples were collected from September to December 2022 using High Volume Air Samplers (TE-6070VFC+X-2.5). The PM_2.5_-containing filter membranes were wrapped in aluminum foil and stored at 4 °C to preserve sample integrity after PM_2.5_ collection. The PM_2.5_ samples were extracted from the filters following the established protocol ^25,26^. In brief, the filter membranes were cut into small pieces and sonicated in deionized water to isolate the water-soluble fraction. After sonication, the filter pieces were removed and rinsed with water to collect any residual particles. The resulting aqueous extract was frozen at −20□°C for 24 hours and then lyophilized. The same filter pieces were subsequently subjected to a second round of ultrasonic oscillation in methanol to isolate the methanol-soluble fraction. This methanol extract was then concentrated to dryness using a gentle nitrogen blow. The dried powders from both the water and methanol fractions were combined and stored at −20□°C until use. The PM_2.5_ powders were reconstituted in ALF to prepare a uniform suspension at a concentration of 2.5 µg/µL ^27^.

### Behavioral tests

A series of behavioral tests was conducted on the day following the final treatment to assess memory performance. The mice underwent one test per day, with assessments carried out between 10:00 am and 6:00 pm.

### Open field test (OF)

For the OFT, mice were habituated to dim light in the testing room for 2 h before testing, as previously described ^28,29^. Mice were then allowed to explore freely for 10 minutes in an acrylic box (dimensions 40 cm × 40 cm × 40 cm), and their behavior was recorded. The first 5 minutes of the recording were analyzed using AnyMaze software (USA) to determine time spent in the center region and distance traveled.

### Novel object recognition task (NOR)

The assessment of recognition memory was performed according to established protocols examining object exploration in mice ^30,31^. During the training, animals were placed in the testing chamber, which contained two identical objects placed 6 cm from the chamber walls. The mice were allowed to freely explore the objects for 10 minutes. Following this initial exploration, the mice had a 2-hour inter-trial interval before being returned to the testing chamber for the testing phase. In the subsequent session, one of the familiar objects was replaced with a novel object that differed in size and appearance, while the other remained unchanged. The time spent exploring both the familiar and novel objects was quantified throughout the 5-minute test period in a sample-blind manner. Three behavioral indices were calculated: the familiar object exploration ratio (familiar/familiar + novel), the novel object exploration ratio (novel/familiar + novel), and the novel object discrimination index (novel − familiar/familiar + novel).

### Y maze test

The Y maze was conducted as previously described ^28^ and in accordance with the established protocol ^32^. The maze consisted of three arms arranged at 120° angles, with each arm measuring 30 cm in length, 8 cm in width, and 6 cm in height. During the habituation phase, mice were placed in the designated start arm and allowed to explore the maze freely for 10 min, during which one arm was blocked by a removable barrier. After a 2 h inter-trial interval, animals were returned to the maze for a 5 min test session, during which the previously obstructed arm was made accessible, allowing exploration of all three arms. Three behavioral measures were subsequently calculated: the familiar arm exploration ratio (time spent in the familiar arm/total time spent in both familiar and novel arms), the novel arm exploration ratio (time spent in the novel arm/total time spent in both familiar and novel arms), and the novel arm discrimination index (difference between time spent in the novel and familiar arms/total time spent in both arms).

### Forced swimming test (FST)

The FST was conducted using the previously described protocol ^29,33^. Individual mice were placed in a transparent acrylic cylindrical container (15□cm in diameter, 30□cm in height) filled with water to a depth that prevents escape or contact with the bottom via the hind limbs or tail. The test typically lasts six minutes, with immobility time quantified during the final four minutes to allow initial escape-driven activity to subside. Behavioral analysis was performed by two trained, sample-blind observers. An increase in total immobility duration is regarded as an indicator of passive coping mechanisms and heightened behavioral despair, which are considered reflective of a depression-like phenotype. Following the test, mice were removed from the water, gently dried with a red-light infrared heat lamp, and returned to their home cages to minimize physiological stress.

### Tissue preparation

Tissues were processed as we previously performed ^34,35^. Mice were administered deep anesthesia with isoflurane prior to the experimental procedures. Animals were perfused with 1x phosphate-buffered saline (PBS), followed by 4% paraformaldehyde in 0.01 M PBS. The brains were isolated and post-fixed in 4% PFA at 4 °C overnight. The brains were stored in 30% sucrose solution at 4 °C until they sank, and coronal brain sections 30 μm thick were prepared using a vibratome (Leica Biosystems, Germany) in a 1-in-6 series. The brain slices were then stored in a cryoprotectant solution (30% glycerol and 30% ethylene glycol) at 4 °C until use.

### Immunohistochemistry

Immunohistochemistry was performed using the free-floating DAB (3,3’-diaminobenzidine) method according to our established protocol ^34^. Antigens were retrieved using the citric acid heat retrieval method. After three washes in 1x PBS for 10 minutes each, brain slices were incubated overnight at room temperature with anti-ki67 (1:200, Cell Signaling Technology) and rabbit anti-doublecortin (DCX) (1:1000, Cell Signaling Technology). The next day, slices were incubated with secondary antibodies, biotinylated goat anti-mouse or goat anti-rabbit IgG (1:200; Vector Laboratories, Burlingame, CA, USA), for two hours at room temperature. Following three washes for 10 minutes each with PBS, visualization of positive cells was achieved using the VECTASTAIN ABC kit (HRP) and the DAB peroxidase substrate kit for 2 hours, as previously reported ^36^.

### Quantification of proliferating cells and immature neurons

Quantification of Ki67^+^ and DCX^+^ Cells was performed by a trained researcher using a Nikon series Eclipse H600L microscope. The total number of labeled cells located in the hippocampal dentate gyrus (DG) was counted in a sample-blinded manner, adapting a previously established protocol ^37,38^. Ki67-positive and DCX-positive cells were quantified across the dorsal DG (from Bregma −1.34 to −2.54 mm) and ventral DG (from Bregma −2.54 to −3.80 mm). Only cells located within the subgranular zone (SGZ) were included in the analysis, as performed previously ^36,38^. The total number of immunopositive cells was estimated by multiplying the average number of labeled cells per section by the total number of 30-μm-thick sections encompassing the quantified DG regions, adapting a previous protocol ^34,36^.

### Brain slice preparation for electrophysiology

Brain slices were prepared following the established protocol ^39^. Adult mice, aged 8-9 weeks, were deeply anesthetized with isoflurane and quickly decapitated. The brains were rapidly immersed in ice-cold, oxygenated (95% O_2_/5% CO_2_) artificial cerebrospinal fluid (aCSF) composed of 125 mM NaCl, 2.5 mM KCl, 1.25 mM NaHPO_4_, 25 mM NaHCO_3_, 2 mM CaCl_2_, 1.3 mM MgCl_2_, and 10 mM dextrose, pH 7.3. Coronal hippocampal sections (350 µm thick) were cut using a vibratome (Leica, VT2000S) in well-oxygenated, ice-cold aCSF. The slices were then gently transferred to a chamber and continuously incubated in oxygenated aCSF at 32°C for 1 hour to facilitate recovery before recording.

### Measurement of synaptic plasticity

Field excitatory postsynaptic potentials (fEPSPs) were recorded from the dentate gyrus (DG) using a multielectrode array system (Alpha MED Scientific Inc., Japan), as previously described ^39^. Recording electrodes on the probe (P515A, Alpha MED Scientific Inc., Osaka, Japan) were positioned in the middle molecular layer of the suprapyramidal blade of the DG to stimulate and record evoked responses. Slices were continuously perfused with oxygenated artificial cerebrospinal fluid (aCSF) at ∼2 mL/min. Signals were acquired using MED-A64MD1 and MED-A64HE1S amplifiers and Mobius software. For each slice, stimulation intensity (20–30 μA) was adjusted to evoke an fEPSP slope equal to 40–50% of the maximal response without eliciting population spikes. Baseline responses were recorded every 15 s for at least 20 min. LTP was then induced by high-frequency stimulation consisting of four trains of 50 pulses at 100 Hz separated by 30 s, delivered in the presence of picrotoxin to block GABA receptor–mediated inhibition. Input-output curves were generated by increasing stimulus intensity in 10-μA increments, with pulse duration varied from 30 to 300 μs in 30-μs steps. Paired-pulse responses were recorded using five paired stimuli delivered at a 50-ms inter-pulse interval with 20-s intervals between pairs. Electrophysiological data were expressed as the percentage of baseline fEPSP slope, calculated from the mean response during the final 20 min before LTP induction.

### Genomic DNA extraction

Fecal samples were collected from male and female C57BL/6J mice (n=6) subjected to chronic PM_2.5_ exposure, as well as from control mice at P57, and stored at −80°C until further processing. For genomic DNA extraction, 180–220 mg of each stool sample was used. Genomic DNA was isolated using the TIANamp Stool DNA Kit (Cat. no. GDP328, TIANGEN Biotech, Beijing, China) according to the manufacturer’s instructions. Library preparation and sequencing were performed by BGI (Beijing, China), generating 150-bp paired-end reads according to standard protocols. DNA fragments underwent end repair, A-tailing, and ligation to full-length Illumina adapters, followed by PCR amplification. All PCR products were purified using Agencourt AMPure XP beads, dissolved in Elution Buffer, and labeled to complete library construction. The size and concentration of the libraries were assessed using an Agilent 2100 Bioanalyzer. Libraries that met quality criteria were sequenced on the HiSeq platform, with each library sequenced to its specified insert size.

### 16S rRNA gene amplicon sequencing and bioinformatics

Genomic DNA was extracted using the PowerSoil Pro kit (Qiagen, USA), and the 16S rRNA gene V4 hypervariable region was amplified using 515f/806r primers ^40^. Library size and concentration were measured using the Agilent 2100 Bioanalyzer. Quality filtering of the raw reads was performed using fastp (v.0.23.1) with default settings ^40^, and chimeric sequences were removed using the UCHIME algorithm ^41^. On average, 64000 ± 8,911 high-quality reads were obtained per sample. Amplicon sequence variants (ASVs) were generated and denoised using the DADA2 plugin ^42^ in QIIME2 (v.2023.07) ^43^. Taxonomic assignment of ASVs was performed using the SILVA 138 database (99% similarity) ^44^. Alpha-diversity metrics, including ASV Shannon and Simpson indices, were calculated using the q2-diversity plugin in QIIME2 after rarefying the samples to a depth of 34,600 reads. Differences in alpha diversity across different treatments and sex were analyzed using two-way ANOVA with Tukey’s post hoc test.

### Metagenomic sequencing

Library preparation and sequencing were performed by Novogene (Beijing, China), generating 150-bp paired-end reads according to the manufacturer’s protocol. Briefly, extracted DNA was fragmented to an average size of 350 bp using the Covaris LE220R-plus system (Woburn, MA, USA). The fragments were then end-polished, A-tailed, and ligated with full-length Illumina sequencing adapters, followed by PCR amplification using the Nextera DNA Flex Library Preparation Kit (Illumina, San Diego, CA, USA). The resulting PCR libraries were quantified, pooled, and sequenced on a NovaSeq 6000 platform (Illumina, San Diego, CA, USA). The raw sequence data were first processed to remove sequence adapters using AdapterRemoval (v.2.3.3) ^45^. Next, quality filtering and contaminant decontamination against both human and mouse were performed using KneadData (v.0.10.0), utilizing the human genome hg37 as the reference and the default parameters. Finally, unpaired reads were removed from the paired-end FastQ files using fastq-pair (v.1.0) ^46^, yielding an average of ∼37.9 (SD ± 15.8) million paired-end reads retained per dust sample for downstream analysis.

### Analysis of bacterial diversity, composition, and function

Taxonomic classification of the paired-end reads was performed using Kraken2 (v.2.1.2) ^47^ with the standard Kraken2 database (v.2024.1.12), and species-level abundance was estimated using Bracken (v.2.9). Species-level taxonomy was used to identify factors associated with community diversity and compositional changes. HUMAnN3 (v.3.7) ^48^ was used to profile the potential metabolic functions of the metagenomes, including the analysis of KO assignments and metabolic pathways using MetaCyc v24.0. Principal coordinate analysis based on Bray–Curtis dissimilarity was performed using the “vegdist” function in the R package “vegan” (v.2.6-4). PERMANOVA with Benjamini–Hochberg adjustment was applied using the “adonis2” function in “vegan” with 999 permutations to test the influence of treatment and gender on microbiome composition. The associations between functional data and treatment were determined using MaAsLin2 ^49^ with the generalized linear model. A *p*-value ≤□0.05 was considered statistically significant. Statistical significance between two groups was assessed using the Wilcoxon rank-sum test (WRST) with false discovery rate adjustment via the Benjamini–Hochberg method. Comparisons among more than two groups were analyzed with the Kruskal–Wallis rank-sum test. Both tests were performed in the R package “stats” (v.4.2.2). A *p*-value of ≤ 0.05 was considered statistically significant.

Contigs with lengths > 1,000□bp were binned into metagenome-assembled genomes (MAGs) using the “binning” function of MetaWRAP (v.1.3) ^50^. The resulting MAGs were further refined using the “bin_refinement” function of MetaWRAP with the parameter “-c 50 -× 10” and dereplicated using the “dRep dereplicate” function of dRep (v.3.2.2). In total, 190 representative MAGs (rMAGs) with contamination ≤ 10% and completeness ≥ 50% were generated. The taxonomy of the 190 rMAGs was annotated using GTDB-TK (v.2.1.0). The phylogeny of the MAGs was performed using PhyloPhlAn3 ^51^ and visualized using the Interactive Tree of Life (iTOL) tool (https://itol.embl.de). The open reading frames (ORFs) of the rMAGs were predicted using Prodigal (v.2.6.3) with the parameter “-p meta,” and the annotation of ORFs was conducted using eggnog-mapper (v.2.1.13) ^52^.

### Synaptoneurosome and total protein extraction

For synaptoneurosome extraction, the mice were euthanized immediately after PM_2.5_ exposure. The hippocampi were dissected and homogenized according to the manufacturer’s protocol in the ice-cold Syn-PER synaptic protein extraction reagent kit (Thermo Fisher Scientific, USA). The hippocampal homogenates were centrifuged at 1200× *g* at 4 °C for 10 min to collect the cytosolic fraction, which was then further centrifuged at 15,000× *g* at 4 °C for 20 min. The resulting synaptosomal fraction was resuspended in the Syn-PER reagent. For total protein extraction, hippocampi were homogenized on ice in radioimmunoprecipitation assay (RIPA) buffer (Abcam, Cambridge, UK) supplemented with Halt™ phosphatase/protease inhibitor cocktail. The homogenates were sonicated (20-second pulse at 50% duty cycle) and then centrifuged at 14,000 × g for 30 minutes at 4°C, following previously established protocol ^29^. The resulting supernatant was collected, and total protein concentration was determined via a Bradford assay (Bio-Rad Laboratories, Hercules, CA, USA). All samples were aliquoted and stored at –80°C for subsequent analysis.

### Western blotting

Samples were diluted in a solution containing Syn-PER reagent or RIPA buffer, 4% 1× Laemmli buffer (Bio-Rad Laboratories, USA), and 5% β-mercaptoethanol (Abcam, USA), then linearized at 95 °C for 10 min. Each lane was loaded with 30 μg of homogenate protein and separated on an 8% SDS-polyacrylamide gel (Bio-Rad Laboratories, USA), then transferred to polyvinylidene fluoride (PVDF) membranes (Bio-Rad Laboratories, USA). Five percent bovine serum albumin (Sigma, USA) in Tris-buffered saline with Tween-20 was used to block nonspecific protein targets for 1 hour at room temperature. The membranes were incubated overnight with primary antibodies diluted in a blocking solution. The primary antibodies used for the target proteins are listed in the Supplementary Table 1. After three washes in TBST for 10 minutes each, the membranes were incubated with horseradish peroxidase-conjugated secondary antibodies for 1 hour at room temperature. After a final wash with TBST, the bands were visualized using an enhanced chemiluminescent (ECL) detection kit (Santa Cruz Biotechnology, Inc., USA) and documented using a transilluminator (Bio-Rad Laboratories, USA). The resulting bands were analyzed using ImageJ.

### Oxidative stress markers measurement

Following the deep anesthesia with isoflurane, mice were euthanized via decapitation. Whole blood samples were immediately collected from the decapitation site into 1.5 ml Eppendorf tubes. Post-collection, samples were left undisturbed at room temperature for 30 minutes to facilitate complete clot formation. Subsequently, serum was separated by centrifugation at 1,000 × g for 20 minutes at 4°C ^53^. The serum aliquots were then carefully stored at −80°C until further analysis. Immediately after blood collection, the hippocampi were rapidly dissected on ice to preserve tissue integrity and prevent protein degradation. Dissected tissues were homogenized in an appropriate lysis buffer with a mechanical homogenizer, then sonicated at 50% amplitude for 20 seconds to ensure complete cell lysis. Homogenates were centrifuged at 14,000 × g for 30 minutes at 4°C to pellet cellular debris, and the supernatants were collected and stored at −80°C until subsequent biochemical assays. Quantification of oxidative stress biomarkers in both serum and hippocampal tissue was performed using commercially available enzyme-linked immunosorbent assay (ELISA) kits according to the manufacturers’ protocols. Glutathione peroxidase (GSH-Px) levels were measured using a mouse-specific ELISA kit (Catalog No: CSB-E13068m; Cusabio, China). Superoxide dismutase (SOD) concentrations were determined with the Mouse Superoxide Dismutase ELISA Kit (Catalog No: RK03201; ABclonal, China). Lipid peroxidation was assessed by measuring malondialdehyde (MDA) levels using an established assay kit (Catalog No: A003-1-1; Nanjing Jiancheng Bioengineering Institute, China). All samples were assayed in duplicate, with concentrations derived from standard curves generated for each assay.

### Intrahippocampal injection for PERK knockdown

AAV2/9 carrying shRNA targeting PERK (5.00 × 10¹² vg/mL) was obtained from BrainVTA and stored in aliquots at −80□°C. Mice were anesthetized using a mixture of ketamine (100□mg/kg) and xylazine (10□mg/kg), then positioned in a stereotaxic apparatus (RWD, Shenzhen, China). After local sterilization and scalp incision, bilateral craniotomies were performed using a high-speed microdrill ^54^. A glass micropipette connected to an ultra-micro injection pump (Nanoliter2010, WPI, USA) and controlled via an injection controller (Micro4, WPI, USA) was used to deliver the AAV solution into the hippocampal CA1 region at the following coordinates: AP −2.0□mm, ML ±1.5□mm, DV −1.5 to −0.6□mm ^55^. The micropipette was left in place for 3 minutes post-injection to allow for proper diffusion. Mice were allowed to recover for three weeks following surgery (Fig. S6b-d).

### AAV8-TBG-Cre–Mediated Hepatocyte-Specific Deletion of FMO3

FMO3 floxed mice (FMO3*^flox/flox^*) were kindly provided by Prof. Kenneth Cheng (Department of Health Technology and Informatics, The Hong Kong Polytechnic University) ^56^. To achieve hepatocyte-specific deletion of FMO3, we used an adeno-associated virus serotype 8 vector encoding Cre recombinase under the control of the thyroxine-binding globulin (TBG) promoter (AAV8-TBG-Cre; WZ Biosciences Inc., China), which drives liver-selective transgene expression. Following established protocols (Sun et al., 2024) ^57^, 5-week-old FMO3*^flox/flox^* mice received 1 × 10¹¹ vector genomes (vg) per mouse via lateral tail vein injection. Immediately prior to administration, 10 μL of viral stock (1 × 10¹³ vg/mL) was diluted in sterile phosphate-buffered saline (PBS) to a final volume of 100 μL and injected using a 27-gauge syringe. Control mice received an equivalent dose and volume of AAV8-TBG-GFP to control for potential vector-associated immunogenicity or hepatotoxicity. Mice were maintained for 3–4 weeks after injection to allow efficient Cre-mediated genomic recombination and turnover of pre-existing FMO3 protein (Fig S5a & b).

### Statistical analysis

Sample sizes were determined based on effect sizes observed in prior studies/lab experience. Animals were randomly assigned to treatment groups, and all data were analyzed without exclusion. Outcome assessments were performed by investigators blinded to the treatment group. Complete statistical reporting, including test justifications and group variations, is included in the figure legends.

The data were presented as mean ± standard error of the mean (SEM). Paired t-tests were conducted to compare the exploration ratio to the novel and familiar arms in the Y-maze task and NOR. Two-way ANOVA and Tukey Post hoc tests were performed for multiple comparisons. Statistical analyses were performed using Prism 8.0.1 (GraphPad Software, USA). A P < 0.05 was considered statistically significant.

## Results

### Chronic PM_2.5_ exposure impairs recognition and spatial working memory in mice

To determine whether PM_2.5_-induced cognitive impairment could be attributed to nonspecific changes in locomotor capacity or general health status, mice were first evaluated in the OFT. Total distance travelled and traveling speed were comparable between control and PM_2.5_-exposed mice in both sexes (Fig. 1b-c). In contrast, PM_2.5_ exposure elicited sex-specific effects on reducing time spent in the center in females but not males, indicating PM_2.5_ increased anxiety-like behavior in females only (Fig. 1d).

We next assessed recognition memory using the NOR task. Control mice of both sexes showed a robust preference for the novel object, consistent with intact recognition memory (Fig. 1e, f). This preference was abolished following PM_2.5_ exposure, with exposed mice exhibiting a shift toward exploration of the familiar object, indicative of impaired recognition memory (Fig. 1e, f). Two-way ANOVA revealed a highly significant main effect of treatment, with no effect of sex and no sex × treatment interaction (Fig. 1d), indicating that PM_2.5_ disrupted recognition memory similarly in males and females. Post hoc analyses confirmed a significant reduction in the NOR index in PM_2.5_-exposed mice of both sexes compared to their sex-matched controls (Fig. 1g).

Similar results were also observed in the Y-maze task, which assesses spatial working memory. Control mice preferentially explored the novel arm, consistent with intact spatial memory performance (Fig. 1h, i). In contrast, this novelty preference was lost in PM_2.5_-exposed mice, indicating a deficit in spatial working memory (Fig. 1h, i). Although PM_2.5_-exposed females showed a significant bias toward the familiar arm in the exploration ratio analysis (Fig. 1i), two-way ANOVA revealed a significant main effect of treatment, with no significant effect of sex or sex × treatment interaction (Fig. 1j). Post-hoc comparisons further demonstrated that PM_2.5_ significantly reduced novel arm exploration in both males and females (Fig. 1j).

Together, these data show that chronic PM_2.5_ exposure did not affect locomotor activity but impaired mood and cognitive function. PM_2.5_ impairs memory performance in both sexes, whereas the anxiety-like response appears to be more pronounced in females.

### Chronic PM_2.5_ exposure impairs hippocampal structural and functional plasticity

To determine whether chronic PM_2.5_-induced behavioral deficits are linked to hippocampal impairment, we first examined changes in hippocampal structural plasticity. We quantified doublecortin-positive (DCX+) immature neurons and Ki67-positive (Ki67+) proliferating cells in the dorsal and ventral dentate gyrus (DG) (Fig. S1). Across the entire dentate gyrus, two□way ANOVA revealed a significant main effect of treatment on the number of DCX+ immature neurons, with no significant main effect of sex and no sex × treatment interaction (Fig. 2b), indicating an overall reduction in immature neuron number following PM_2.5_ exposure. Although sex□stratified post hoc tests revealed a significant decrease in DCX+ cells in females but not in males, this sex-specific pattern is exploratory. Total Ki67+ cell counts revealed a significant sex × treatment interaction (Fig. 2d), indicating that the suppressive effects of PM_2.5_ on cell proliferation in a sex-dependent manner. Post hoc test confirmed a marked reduction in proliferating cells in females, whereas no significant effect was detected in males. These findings indicate that chronic PM_2.5_ exposure disrupts adult hippocampal structural plasticity, with female mice showing heightened vulnerability. These data further suggest that impaired hippocampal structural plasticity may represent a cellular substrate contributing to the behavioral deficits induced by chronic PM_2.5_ exposure.

**Fig. 2.**
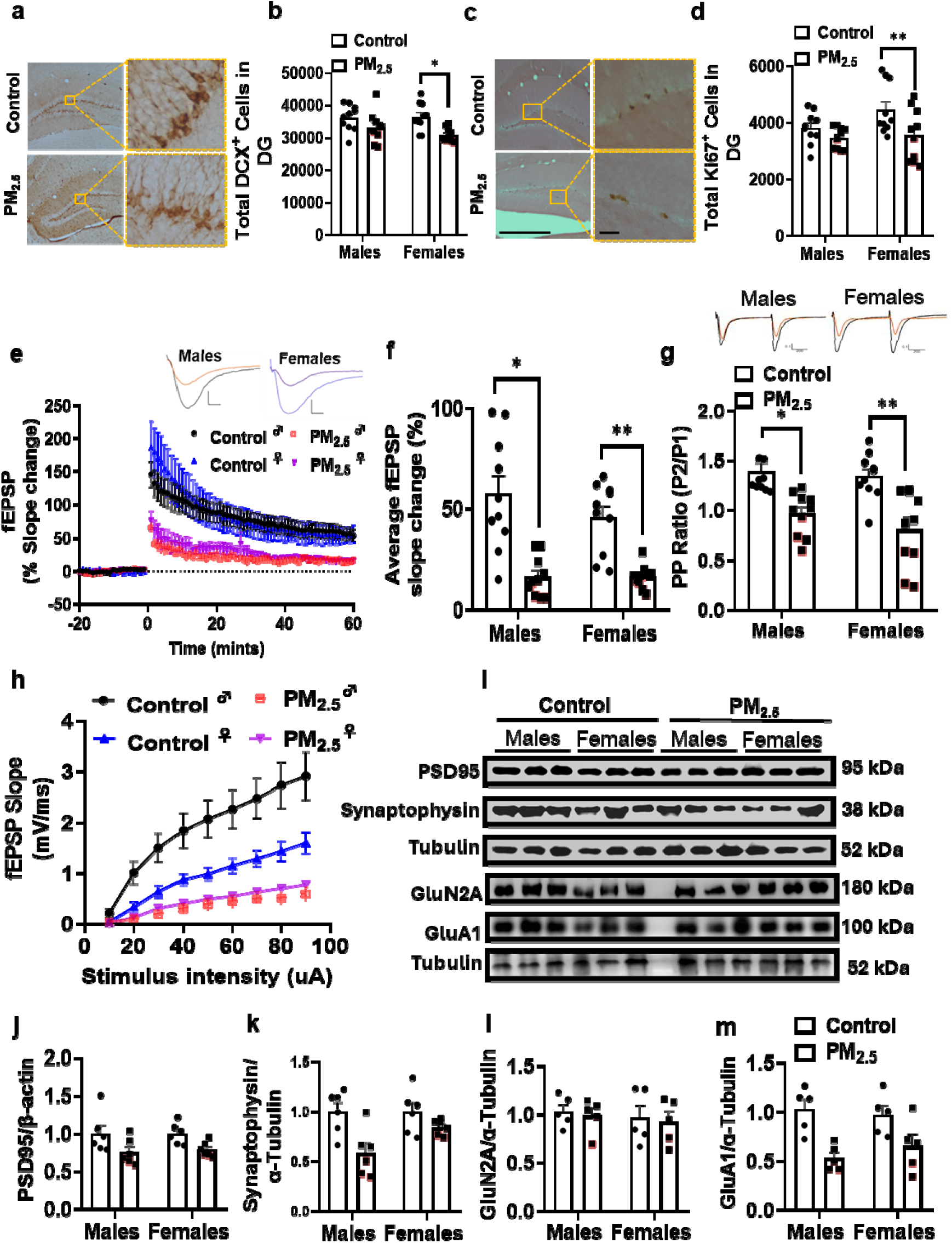
Chronic PM_2.5_ exposure impairs hippocampal structural and functional plasticity. a-b. Representative immunohistochemical images and quantification of (**a, b**) DCX+ immature neurons and (**c, d**) Ki67+ proliferating cells in the hippocampal dentate gyrus (DG). PM_2.5_ reduces cell proliferation primarily in females. **b** DCX+ cell quantification showed a significant main effect of treatment [F(1,9)=9.95, P=0.012]. Post hoc Tukey tests confirmed significant reductions in females (P=0.0191) but not in males (P=0.0891). **d** Ki67+ counts showed a significant sex × treatment interaction [F(1,9)=8.00, P=0.020] and main treatment effect [F(1,9)=12.45, P=0.006]. Two-way ANOVA with Post hoc Tukey tests revealed that PM_2.5_ decreased cell proliferation in females (P=0.0041) but not in males (P=0.3865). n = 10 mice/group. **e-h** Electrophysiological recordings: PM_2.5_ impairs hippocampal LTP and synaptic efficacy in both sexes. **f** Average fEPSP slopes (50–60 min post-HFS) showed a significant main effect of treatment [F(1,9)=51.99, P<0.0001]. Post hoc Tukey tests confirmed impaired LTP in both males (P=0.0260) and females (P=0.0018). **g** Paired-pulse ratio (PPR) analysis showed a significant treatment effect [F(1,9)=45.04, P<0.0001]. Post hoc Tukey tests indicated impaired short-term plasticity in both males (P=0.0025) and females (P=0.0359). **h** Input–output (I/O) curves demonstrated reduced synaptic efficacy post-LTP induction in both sexes (P<0.0001). n = 10 slices from 5 mice per group. **i-m** Western blot analysis of hippocampal synaptoneurosomes: Representative western images (**i**). Two-way ANOVA revealed significant main effects of treatment for (**j**) PSD95 [F(1,5)=9.992, P=0.0251], (**k**) synaptophysin [F(1,5)=21.40, P=0.0057], and (**m**) GluA1 [F(1,4)=12.39, P=0.0245], while (**l**) GluN2A remained unchanged [F(1,4)=0.1216, P=0.7448]. No significant sex effects or sex × treatment interactions were observed for any protein (all P>0.10). Post hoc Tukey tests for PSD95 (males P=0.3927, females P=0.1317), synaptophysin (males P=0.1698, females P=0.3184), and GluA1 (males P=0.076, females P=0.218) showed consistent trends across both sexes. n=10/group. All data are expressed as mean ± SEM.

To test whether chronic PM_2.5_ exposure disrupts hippocampal synaptic function, we assessed electrophysiological recording in the DG. High-frequency stimulation of the perforant path induced robust LTP in control slices, whereas this response was markedly attenuated in slices from PM_2.5_-exposed mice (Fig. 2e). Quantification of the potentiated response 50 to 60 min after stimulation revealed a significant main effect of treatment, and post hoc analyses confirmed a reduction in LTP formation in both males and females (Fig. 2f). Although the deficit appeared more pronounced in females, the absence of a significant sex × treatment interaction indicates that PM_2.5_-induced impairment of LTP was not statistically sex dependent. This deficit in activity-dependent synaptic strengthening was accompanied by impairments in DG synaptic function in both sexes. Input-output analysis showed reduced synaptic efficacy in PM_2.5_-exposed mice following LTP induction in both sexes (Fig. 2h). In parallel, paired-pulse ratio (PPR) measurements revealed a significant effect of treatment, with post hoc testing confirming altered presynaptic function in both males and females (Fig. 2g). Together, these findings indicate that chronic PM_2.5_ exposure compromises both synaptic function and presynaptic release in the DG.

To identify molecular correlates of the synaptic deficits induced by chronic PM_2.5_ exposure, we next quantified the expression of key pre- and postsynaptic proteins in the hippocampus by western blotting. Two-way ANOVA revealed a significant main effect of treatment on the postsynaptic scaffolding protein PSD95, with no significant main effect of sex or sex × treatment interaction (Fig. 2j), indicating an overall reduction in postsynaptic structural integrity following PM_2.5_ exposure (Fig S2a). A similar pattern was observed for the presynaptic vesicle protein synaptophysin, which was also significantly reduced by treatment in the absence of significant sex effects or interaction (Fig. 2k; S2b). In parallel, expression of the AMPA receptor subunit GluA1 was significantly decreased in PM_2.5_-exposed mice, again with no effect of sex and no sex × treatment interaction (Fig. 2m; S2d), consistent with impaired excitatory synaptic transmission. By contrast, levels of the NMDA receptor subunit GluN2A were unchanged (Fig. 2l; S2c). Although post hoc comparisons within each sex did not reach statistical significance for PSD95, synaptophysin, or GluA1, the significant treatment effects detected by two-way ANOVA indicate a consistent downregulation of synaptic proteins across sexes. Together, these molecular changes align with the electrophysiological evidence of impaired dentate gyrus plasticity and support the conclusion that chronic PM_2.5_ exposure disrupts synaptic plasticity required for normal hippocampal synaptic function.

### Chronic PM_2.5_ exposure alters gut microbiome and elevates TMAO levels

Given the emerging role of the gut microbiota in regulating brain function and cognition through the gut–brain axis, we next asked whether chronic PM_2.5_ exposure induces microbial alterations that could contribute to the observed neurobehavioral and hippocampal deficits.

To determine the gut microbial response to chronic PM_2.5_ exposure, we profiled fecal communities by 16S rRNA gene amplicon sequencing and compared microbial diversity and community structure between control and exposed mice (Fig. 3a, b). In females, PM_2.5_ exposure significantly reduced bacterial species diversity while increasing community evenness, indicating a marked shift in alpha-diversity; by contrast, no significant alpha-diversity changes were detected in males (Fig. 3a). Despite this sex difference in within-sample diversity, principal coordinate analysis demonstrated a clear separation between control and PM_2.5_-exposed microbiomes independent of sex, and PERMANOVA confirmed a significant treatment effect on overall community composition (R² = 0.15, P = 0.002) (Fig. 3b, c). We next examined taxonomic features previously implicated in cognitive regulation and neurobehavioral function ^58–60^. PM_2.5_ exposure significantly enriched the phylum Bacillota, the family Bifidobacteriaceae, and the genus *Bifidobacterium* relative to controls (adjusted P < 0.05; Fig. 3d). In addition, *Edwardsiella tarda* was significantly increased in PM_2.5_-exposed mice (adjusted P < 0.05), indicating selective expansion of taxa linked to altered host metabolic and inflammatory states.

**Fig. 3.**
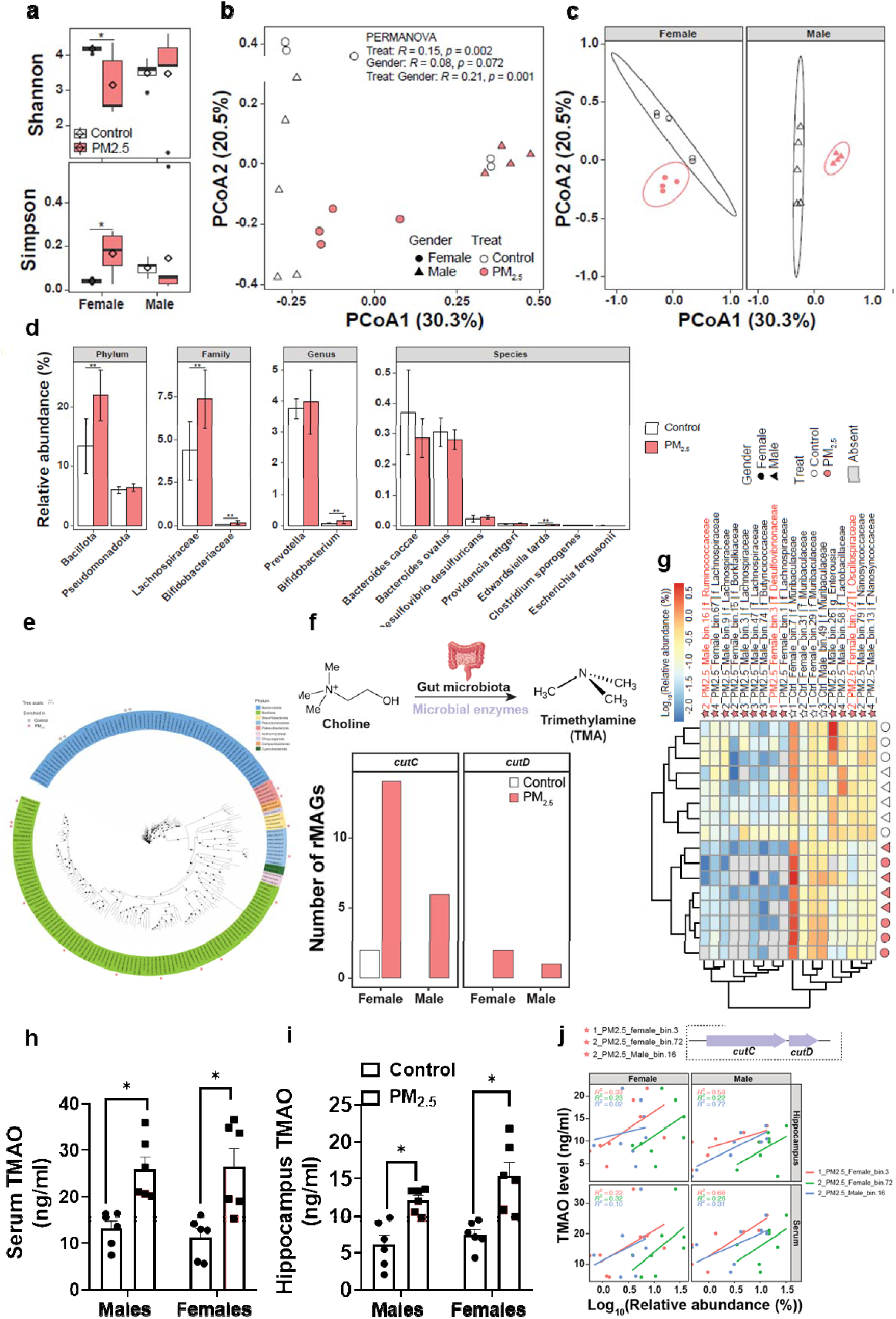
PM_2.5_ induces gut dysbiosis and TMAO production. a-c. Microbial diversity and community structure: PM_2.5_ exposure shifts community composition without significantly altering alpha diversity. **a** Bacterial alpha diversity (Shannon and Simpson indices). **b**, **c** Principal coordinates analysis (PCoA) of Bray–Curtis distances across (**b**) all samples and (**c**) stratified by sex. **d**-**g** Taxonomic and functional profiling of the gut microbiome: PM_2.5_ alters microbial abundance and enriches taxa associated with choline metabolism. **d** Taxonomic composition at the phylum, family, genus, and species levels. **e** Phylogenetic distribution and MAG enrichment analysis. **f** Schematic of microbial choline metabolism and abundance of representative MAGs (rMAGs) harboring cutC/D genes. **g** Heatmap of differentially abundant MAGs (log10 relative abundance). Control n=5/group, PM_2.5_ n=4/group. **h, i** TMAO concentrations in (**h**) serum and (**i**) hippocampus: PM_2.5_ increases TMAO levels in both sexes. **h** Serum TMAO levels showed no significant main effect of sex [F(1,5)=0.1141, P=0.7493] and no sex × treatment interaction [F(1,5)=0.1689, P=0.6981], but there was a significant main effect of treatment [F(1,5)=47.08, P=0.0010]. Post hoc Tukey tests confirmed that PM_2.5_ significantly increased serum TMAO levels in both males (*P=0.0434) and females (*P=0.0476) compared with their respective control groups. **i** Hippocampal TMAO levels similarly showed no significant main effect of sex [F(1,5)=1.891, P=0.2275] and no sex × treatment interaction [F(1,5)=1.164, P=0.3298], but a significant main effect of treatment [F(1,5)=29.36, P=0.0029]. Post hoc Tukey tests revealed that PM_2.5_ significantly increased hippocampal TMAO levels in both males (*P=0.0288) and females (*P=0.0206) relative to controls. **j** A positive trend in Pearson correlations between the relative abundance of three MAGs and serum TMAO levels in both females (r=0.38, P=0.074) and males (r=0.44, P=0.054). n=6/group. All data are expressed as mean ± SEM.

To determine whether these compositional changes were accompanied by altered microbial functional capacity, we performed shotgun metagenomic sequencing. Differential pathway analysis identified 54 MetaCyc pathways that differed between groups, of which 39 were enriched in PM_2.5_-exposed mice (Fig. S3a-c), indicating a broad functional restructuring of the gut microbiome following pollutant exposure. We then reconstructed high-quality draft genomes to resolve organism-level functional contributors. A total of 190 representative metagenome-assembled genomes (rMAGs) were recovered, the majority of which belonged to the phyla Bacteroidota (n = 66) and Bacillota (n = 98) (Fig. 3e). Among these, 14 rMAGs were enriched in PM_2.5_-exposed mice ^61^, whereas only 4 were enriched in controls (Fig. 3e), consistent with a directional microbiome shift induced by exposure. Notably, four of the 14 PM_2.5_-enriched rMAGs were classified as *Lachnospiraceae*, a taxonomic group previously associated with cognition-related phenotypes.

To prioritize candidate microbial pathways potentially relevant to the observed neurobiological phenotypes, we analyzed untargeted serum metabolomics data from an independent cohort of female PM_2.5_-exposed mice (Fig. S4). This analysis revealed broad metabolic perturbations in response to PM_2.5_ exposure, among which TMAO emerged as one of the metabolites elevated in the exposed group. Because TMAO is a host-microbial co-metabolite derived from gut microbial TMA production and has been implicated in systemic inflammation and neurological dysfunction, these preliminary findings prompted us to interrogate microbial genes involved in choline-to-TMA metabolism in the metagenomic dataset.

Given the emerging link between microbiota-derived TMA metabolism and neurobehavioral dysfunction, we next examined microbial genes involved in choline utilization. rMAGs harboring cutC and/or cutD, which encode key enzymes required for TMA production from choline, were more prevalent in PM_2.5_-exposed mice than in controls, with the higher enrichment was observed in females (Fig. 3f). Moreover, three rMAGs containing the complete cutCD gene cluster were selectively enriched in the PM_2.5_ group (Fig. 3g), suggesting that chronic PM_2.5_ exposure increases the abundance of microbial taxa with the capacity to generate TMA, the precursor of TMAO.

Prompted by these metagenomic findings, we next assessed whether PM_2.5_ exposure alters TMAO levels. We therefore quantitatively measured TMAO in serum and brain from adult male and female mice. Two-way ANOVA revealed a significant main effect of treatment on serum TMAO, with no significant main effect of sex or sex × treatment interaction (Fig. 3h), demonstrating that PM_2.5_ robustly elevates circulating TMAO in both sexes. Post hoc analyses confirmed significant increases in serum TMAO in both male and female PM_2.5_-exposed mice relative to sex-matched controls. A similar pattern was observed in the hippocampi (Fig. 3i), where PM_2.5_ exposure significantly increased TMAO concentrations without significant effects of sex or interaction, indicating that PM_2.5_-induced TMAO accumulation extends to the central nervous system. Although the increase appeared greater in females, this trend did not reach statistical significance. Finally, correlation analysis revealed a marginal positive association between the relative abundance of cutC/cut/D taxa and serum TMAO levels in both females (r = 0.38, P = 0.074) and males (r = 0.44, P = 0.054) (Fig. 3j), consistent with a microbiota-dependent contribution to the elevation of serum TMAO levels.

Together, these data show that chronic PM_2.5_ exposure induces a pronounced restructuring of the gut microbiome at both compositional and functional levels and is accompanied by robust increases in circulating and brain TMAO. The enrichment of cutC/cutD-containing bacterial genomes further supports a mechanistic link between PM_2.5_-induced dysbiosis and the activation of the microbial choline-to-TMAO metabolic axis.

### Pharmacological inhibition of TMAO production mitigates PM_2.5_-induced memory deficits

Given the robust elevation of TMAO in both serum and brain following chronic PM_2.5_ exposure, we next asked whether suppressing TMAO production could attenuate the associated neurobiological impairments. To test this, mice were treated with 1% 3,3-dimethyl-1-butanol (DMB), an inhibitor of microbial TMA formation, via drinking water during PM_2.5_ exposure (Fig. 4a). DMB treatment significantly reduced serum TMAO levels in PM_2.5_-exposed mice of both sexes, with no significant sex × treatment interaction (Fig. 4b), indicating effective systemic suppression of TMAO production. In the hippocampus, treatment produced a significant main effect on TMAO concentration (Fig. 4c). Although the sex × treatment interaction was not significant, post hoc tests indicated that DMB significantly lowered hippocampal TMAO in PM_2.5_-exposed females but not males; these sex-stratified findings are exploratory and interpreted with caution.

**Fig. 4.**
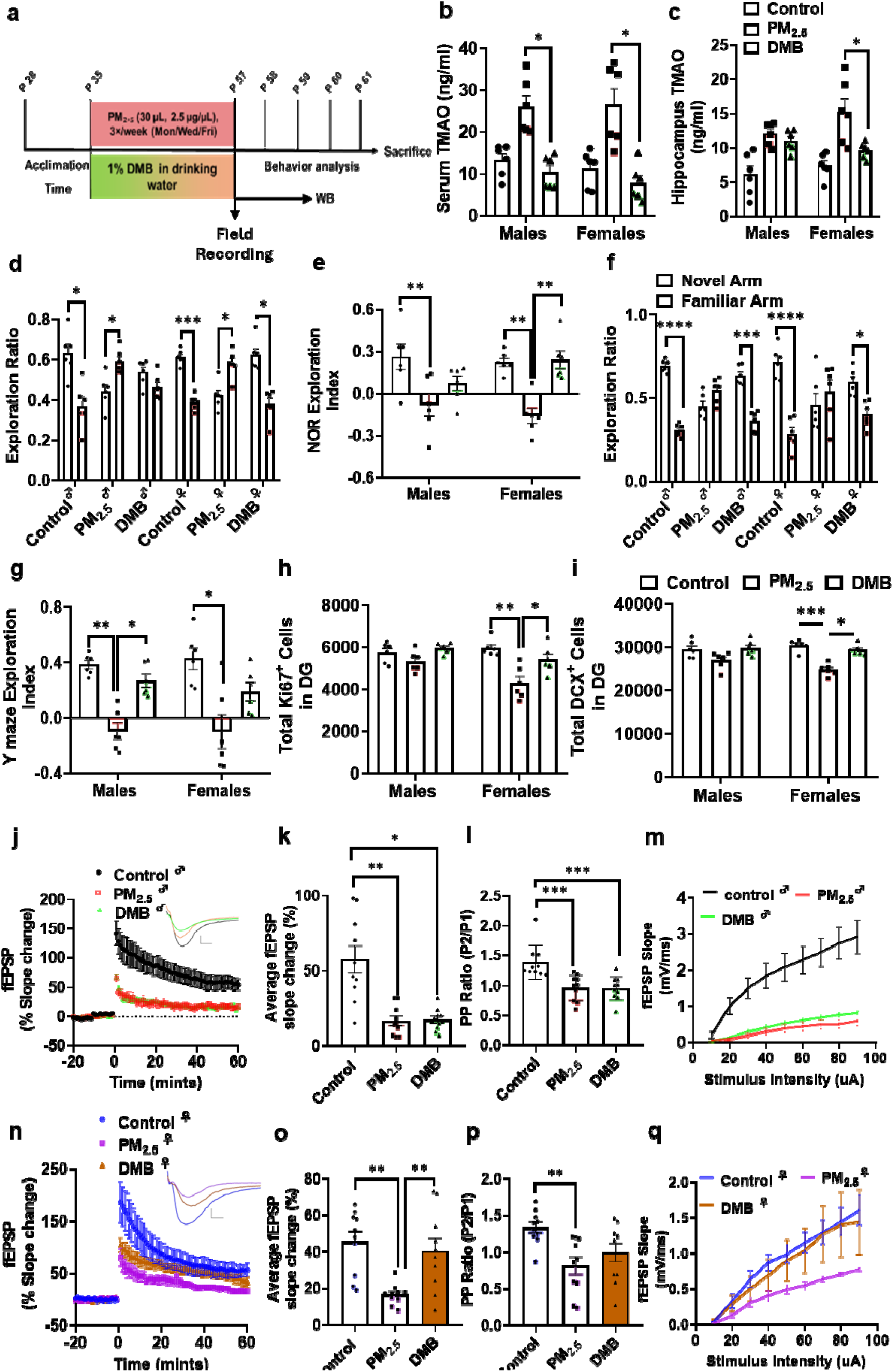
Pharmacological inhibition of TMAO production mitigates chronic PM_2.5_-induced impairments in memory and hippocampal plasticity. **a** Experimental timeline showing the administration of 3,3-dimethyl-1-butanol (DMB) concurrently with PM_2.5_ exposure. **b,c** TMAO concentrations in serum and hippocampus: DMB reduces peripheral TMAO in both sexes but exhibits sex-specific efficacy in the brain. **b** Serum TMAO levels showed a significant main effect of treatment [F(1.219, 6.094) = 23.97, P=0.0021], with no effect of sex (P=0.4358) or sex × treatment interaction (P=0.6607). Post hoc Tukey tests confirmed that DMB significantly reduced PM_2.5_-induced TMAO elevations in both males (P=0.0109) and females (P=0.0454). **c** Hippocampal TMAO levels showed a significant treatment effect [F(1.761, 8.804) = 19.86, P=0.0007] and a near-significant sex × treatment interaction (P=0.0566). Post hoc Tukey tests revealed that DMB significantly reduced hippocampal TMAO in females (P=0.0476) but failed to rescue levels in males (P=0.4712). **d, e** Novel object recognition test (NORT): DMB rescues recognition memory deficits specifically in females. PM_2.5_ abolished novel object preference (**d**: paired t-tests, males *P=0.0458, females *P=0.0352). Two-way ANOVA of the exploration index (**e**): treatment [F(2,30)=17.04, P<0.0001]. Post hoc Tukey tests confirmed significant rescue in females (**P=0.0026) but not in males (P=0.3445). **f, g** Y-maze test: DMB mitigates spatial working memory impairments. PM_2.5_-exposed mice lacked novel arm preference, which was restored by DMB (**f**: paired t-tests, males **P=0.0024, females *P=0.0335). Two-way ANOVA of the exploration index (**g**): treatment [F(2,30)=21.15, P<0.0001]. Post hoc Tukey tests showed a significant rescue in males (*P=0.0180) but not in females (P=0.0911). n=6/group. **h, i** Hippocampal neurogenesis: DMB reverses PM_2.5_-induced reductions in (**h**) Ki67+ proliferating cells and (**i**) DCX+ immature neurons. **h** Two-way ANOVA of Ki67+ counts revealed a significant main effect of treatment [F(1.964, 9.821) = 49.32, P<0.0001] and a sex × treatment interaction [F(1.654, 8.269) = 6.889, P=0.0203]. Post hoc Tukey tests confirmed that DMB significantly rescued proliferation in females (P=0.0351) following PM_2.5_-induced deficits (P=0.0084). **i** Post hoc Tukey tests for DCX+ immature neurons showed that DMB significantly restored neurogenesis in both males (P=0.0322) and females (P=0.0103) compared to PM_2.5_-exposed groups. n=6/group. **j-q** Electrophysiological recordings: DMB restores hippocampal LTP and basal transmission. LTP was significantly rescued in females (**o**: **P=0.0063) but not in males (**k**: P=0.9926). Basal transmission was restored in both sexes (**m, q**: males **P=0.0026, females **P=0.0020). Paired-pulse ratio (PPR) (**l, p**) showed a significant effect of treatment [F(1.67, 15.06)=15.77, P=0.0003; one-way ANOVA followed by post hoc Tukey tests], but no significant DMB rescue. n=10 slices from 5 mice/group. All data are expressed as mean ± SEM.

We next examined whether TMAO inhibition rescues PM_2.5_-induced memory deficits. In the NOR task, control mice of both sexes, as well as DMB-treated PM_2.5_-exposed females, displayed a clear preference for the novel object, whereas PM_2.5_-exposed mice showed impaired recognition memory (Fig. 4d). Two-way ANOVA of the exploration index revealed a significant main effect of treatment, and post hoc analyses confirmed that PM_2.5_ reduced recognition performance in both sexes (Fig. 4e). Notably, DMB restored NOR performance in females but did not significantly improve cognition in males. In the Y-maze, PM_2.5_ exposure abolished preference for the novel arm in both sexes (Fig. 4f), and DMB treatment reinstated this preference. Analysis of the spatial exploration index revealed a significant treatment effect and a significant sex × treatment interaction (Fig. 4g). Post hoc comparisons confirmed that PM_2.5_ impaired spatial exploration in both sexes, whereas DMB rescued this deficit in males but not in females. Thus, inhibition of TMAO production improved PM_2.5_-induced memory dysfunction, although the behavioral rescue showed task- and sex-specific differences.

To determine whether inhibition of TMAO production restores PM_2.5_-impaired hippocampal structural plasticity, we quantified the numbers of proliferating cells and immature neurons in the dentate gyrus. For Ki67+ cells, two-way ANOVA revealed a significant main effect of treatment and a significant sex × treatment interaction, with no significant main effect of sex (Fig. 4h; S5a-d). Post hoc analyses showed that PM_2.5_ significantly reduced Ki67^+^ cell numbers in females, whereas the reduction in males did not reach significance. Notably, DMB treatment significantly rescued the PM_2.5_-induced proliferative deficit in females, while no significant rescue was detected in males. These findings indicate that the effect of TMAO inhibition on adult hippocampal cell proliferation is sex-dependent and more pronounced in females.

A similar rescue was observed for DCX+ immature neurons. Two-way ANOVA showed a significant main effect of treatment and a significant sex × treatment interaction, with no main effect of sex (Fig. 4I; S5e-h). Post hoc comparisons confirmed a significant reduction in DCX+ cell number after PM_2.5_ exposure in females, whereas the decrease in males was not statistically significant. Importantly, DMB treatment significantly increased DCX^+^ cell numbers compared with the PM_2.5_ group in both sexes, indicating recovery of the immature neuronal population. Together, these data show that inhibition of TMAO production reverses PM_2.5_-induced impairments in the DG, with rescued cell proliferation in females and restored immature neurons observed in both sexes.

We next tested whether DMB also alleviates PM_2.5_-induced synaptic dysfunction. In female mice, DMB markedly rescued deficits in LTP and improved basal synaptic transmission (Fig. 4j, k, m, n, o, q). By contrast, the effects on short-term plasticity were more limited. Although paired-pulse facilitation analysis revealed a main effect of treatment (Fig. 4l, p), comparisons between PM_2.5_ and PM_2.5_+DMB groups did not reach significance in either sex, suggesting that TMAO inhibition more effectively restores long-term synaptic plasticity than presynaptic short-term release dynamics.

Together, these data identify the TMA/TMAO pathway as a functionally relevant mediator of PM_2.5_ neurotoxicity. Pharmacological inhibition of TMAO production ameliorated PM_2.5_-induced memory impairment, disrupted hippocampal neurogenesis, and synaptic dysfunction. The extent of rescue varied by sex, with females showing particularly strong recovery in recognition memory, adult neurogenesis, and LTP, whereas spatial memory rescue was more evident in males. These findings support a causal contribution of gut microbiota-dependent TMAO signaling to the hippocampal and behavioral consequences of chronic PM_2.5_ exposure.

### Hepatic FMO3 deficiency reduces TMAO production and prevents PM_2.5_-induced cognitive and hippocampal deficits

To complement the pharmacological studies and further test whether host TMAO biosynthesis contributes to PM_2.5_-induced phenotypes, we used a genetic approach targeting FMO3, the host enzyme that converts microbiota-derived TMA to TMAO (Fig. 5a). Serum TMAO analysis revealed a significant main effect of treatment and a significant sex × treatment interaction, with no main effect of sex (Fig. 5b). In FMO3*^flox/flox^*mice, PM_2.5_ significantly increased serum TMAO in both sexes, whereas this increase was absent in FMO3*^cre/-^* mice. PM_2.5_-exposed FMO3*^cre/-^* mice also had lower serum TMAO than PM_2.5_-exposed FMO3^flox/flox^ littermates in both sexes, indicating that genetic disruption of FMO3 prevents PM_2.5_-induced systemic TMAO accumulation. In the hippocampus, two-way ANOVA revealed a significant main effect of treatment, but no significant effect of sex or sex × treatment interaction (Fig. 5b). Although post hoc comparisons did not reach significance, the overall pattern was similar to that observed in serum, with a trend toward lower hippocampal TMAO in PM_2.5_-exposed FMO3*^cre/-^* females relative to PM_2.5_-exposed FMO3*^flox/flox^* females, this difference was not statistically significant and described as a non-significant trend. These findings indicate that FMO3-dependent metabolism is required for the PM_2.5_-induced increase in systemic TMAO and contributes to the accumulation of TMAO in the hippocampus.

**Fig. 5.**
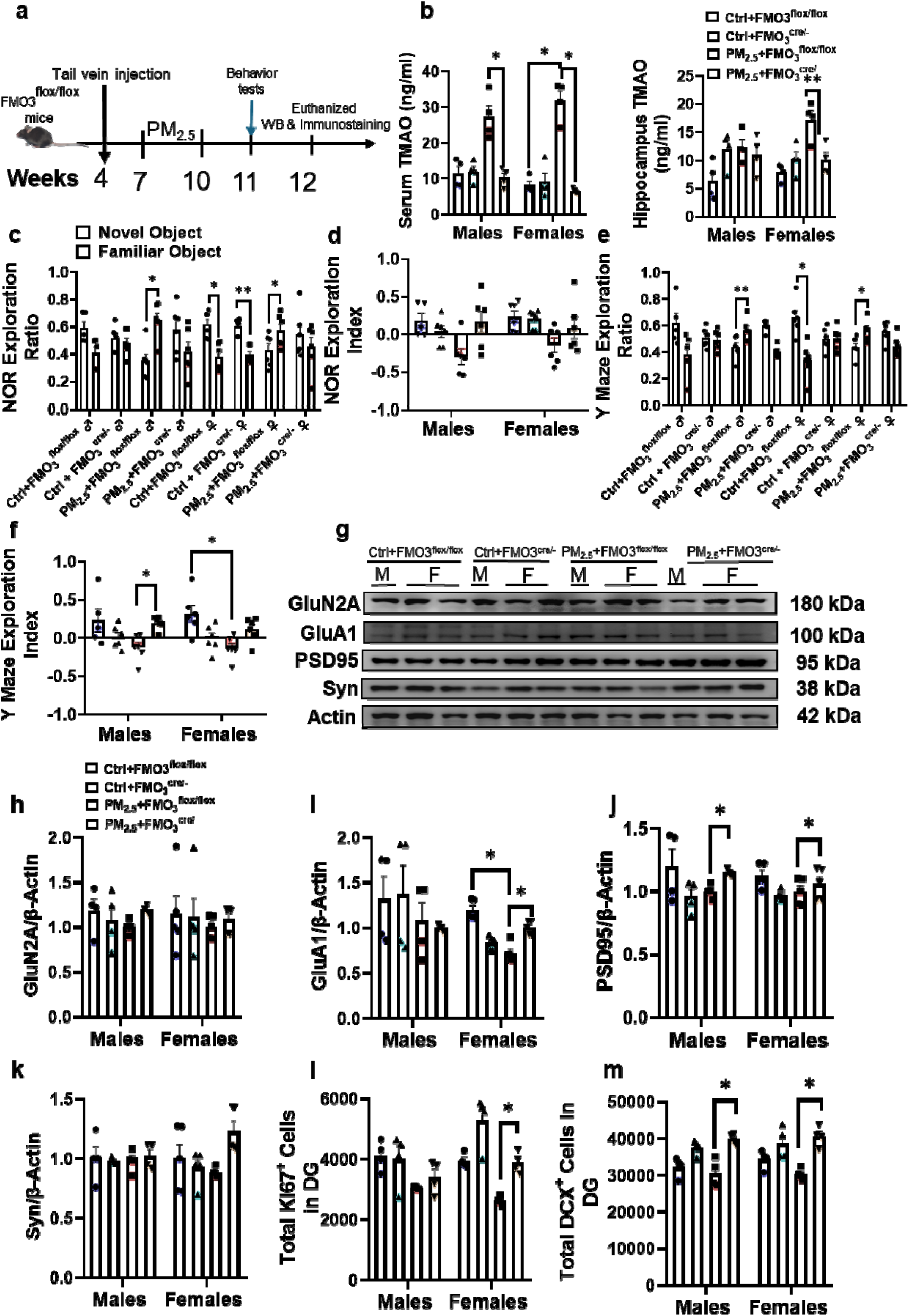
Liver-specific FMO3 deficiency prevents PM_2.5_-induced cognitive and hippocampal dysfunctions. **a** Experimental timeline showing tail vein injection of FMO3-targeted constructs, followed by chronic PM_2.5_ exposure and behavioral assessment. **b** TMAO concentrations in serum and hippocampus: FMO3 deficiency eliminates PM_2.5_-induced TMAO elevation. Serum TMAO showed a significant treatment effect [F(3,20)=42.15, P<0.0001], with post hoc Tukey tests confirming significant reductions in FMO3*^cre/-^* mice compared to FMO3*^flox/flox^* mice under PM_2.5_ exposure (*P<0.05). Hippocampal TMAO levels were significantly increased in PM_2.5_-exposed flox/flox females (**P<0.01) but remained at baseline in cre/- mice. **c, d** Novel object recognition test (NORT): FMO3 deficiency partially preserves novel object preference patterns. PM_2.5_-exposed FMO3*^flox/flox^*mice showed no preference for the novel object (**c**: paired t-tests), whereas FMO3*^cre/-^* mice exhibited restored preference. Two-way ANOVA of the exploration index (**d**) revealed a significant main effect of treatment [F(3,40)=18.24, P<0.0001], with significant rescue observed in both sexes (*P<0.05). **e, f** Y-maze test: PM_2.5_+FMO3^cre/-^ males and Ctrl+FMO3*^flox/flox^* females showed significant preference for the novel arm (paired t-tests; PM_2.5_+FMO3^cre/-^ males **P=0.002, Ctrl+FMO3*^flox/flox^*females *P=0.030). **f** PM_2.5_ exposure reduced the exploration index, while FMO3 deficiency partially preserved performance. Two-way ANOVA revealed a significant main effect of treatment [F(1.388, 6.940)=7.653, P=0.0227], with no significant effect of sex (P=0.9475) or sex × treatment interaction (P=0.6091). Post hoc Tukey tests showed significant differences between PM_2.5_+FMO3^cre/-^ and PM_2.5_+FMO3*^flox/flox^*males (*P=0.0474) and between ALF+FMO3*^flox/flox^* and PM_2.5_+FMO3*^flox/flox^*females (*P=0.0120). n=6/group. Representative images of western blotting (**g). h-m** Synaptic and neurogenesis markers in the hippocampus. **h** GluN2A expression was unchanged across groups, with no significant effects of sex, treatment, or sex × treatment interaction. **i** GluA1 showed no significant main effects, but post hoc analysis identified significant differences in females between ALF+FMO3*^flox/flox^*and PM_2.5_+FMO3*^flox/flox^* (*P=0.0157), ALF+FMO3^flox/flox^ and ALF+FMO3^cre/-^(*P=0.0371), ALF+FMO3*^flox/flox^* and PM_2.5_+FMO3*^cre/-^* (**P=0.0027), and PM_2.5_+FMO3^flox/flox^ and PM_2.5_+FMO3^cre/-^ (*P=0.0364). **j** PSD95 also showed no significant main effects, but post hoc analysis revealed significant differences between PM_2.5_+FMO3*^flox/flox^*and PM_2.5_+FMO3*^cre/-^* in both males (*P=0.0239) and females (**P=0.0093). **k** SYN expression was not significantly altered by sex, treatment, or their interaction. n=4/group. **l, m** Ki67 and Dcx quantification. **l** Ki67 showed significant effects of treatment (P=0.0191) and sex × treatment interaction (P=0.0129); in females, significant differences were detected between ALF+FMO3*^flox/flox^* and PM_2.5_+FMO3*^flox/flox^* (*P=0.0166), PM_2.5_+FMO3*^flox/flox^*and ALF+FMO3^cre/-^ (*P=0.0378), and PM_2.5_+FMO3*^flox/flox^*and PM_2.5_+FMO3^cre/-^(*P=0.0248). **m** DCX showed a significant main effect of treatment (P=0.0016); post hoc analysis revealed significant differences in males between ALF+FMO3*^flox/flox^* and PM_2.5_+FMO3*^cre/-^*(*P=0.0101), PM_2.5_+FMO3*^flox/flox^*and ALF+FMO3*^cre/-^* (*P=0.0472), and PM_2.5_+FMO3*^flox/flox^*and PM_2.5_+FMO3*^cre/-^* (*P=0.0144), and in females between PM_2.5_+FMO3*^flox/flox^* and PM_2.5_+FMO3*^cre/-^*(*P=0.0313). n=4/group. All data are expressed as mean ± SEM.

We next assessed whether genetic suppression of TMAO synthesis protects against PM_2.5_-induced memory deficits. In the NOR task, PM_2.5_-exposed FMO3*^flox/flox^* mice failed to show a preference for the novel object, whereas PM_2.5_-exposed FMO3*^cre/-^* mice retained novel object preference (Fig. 5c). However, the NOR exploration index showed only a significant main effect of treatment, with no significant effect of sex or sex × treatment interaction, and no significant post hoc comparisons within either sex (Fig. 5d). In the Y-maze, control and FMO3-deficient groups generally preferred the novel arm, whereas novelty preference was attenuated in PM_2.5_-exposed FMO3*^flox/flox^* mice (Fig. 5e). The Y-maze exploration index showed a significant main effect of treatment, with no effect of sex and no sex × treatment interaction (Fig. 5f).

Western blot analysis showed no significant differences in GluN2A and synaptophysin (Fig. 5h, k). GluA1 showed no significant effects, although treatment trended toward significance (Fig. 5i). Consequently, sex-stratified post hoc comparisons are exploratory. PM_2.5_-exposed FMO3*^flox/flox^* females tended to have lower GluA1 than control FMO3*^flox/flox^* females, and PM_2.5_-exposed FMO3*^cre/-^*females tended to have higher GluA1 than PM2.5-exposed FMO3*^flox/flox^*females, but these comparisons did not reach significance after correction. Two□way ANOVA revealed no significant main effects or interaction for PSD□95 (Fig. 5j). A post hoc comparison showed higher PSD□95 in PM_2.5_□exposed FMO3*^cre/-^* versus PM_2.5_□exposed FMO3*^flox/flox^* mice, because this comparison was not supported by a significant ANOVA term, treated as exploratory.

Furthermore, we examined changes in adult hippocampal neurogenesis. For Ki67+ cells, two-way ANOVA revealed a significant main effect of treatment and a significant sex × treatment interaction, with no main effect of sex (Fig. 5l). In females, PM_2.5_-exposed FMO3*^flox/flox^* mice had fewer Ki67-positive cells than control FMO3*^flox/flox^* mice and PM_2.5_-exposed FMO3*^cre/-^* mice, indicating protection by FMO3 disruption. No significant pairwise differences were detected in males. DCX analysis also revealed a significant main effect of treatment, but no significant main effect of sex and no sex × treatment interaction (Fig. 5m). Although the exploratory post hoc comparison suggested higher DCX+ counts in PM_2.5_-exposed FMO3^cre/-^ mice relative to FMO3^flox/flox^ littermates in both sexes, these differences were not supported by a significant interaction and are reported as trends. These results indicate that FMO3-dependent TMAO production contributes to PM_2.5_-induced deficits in memory and hippocampal neurogenesis. Genetic disruption of FMO3 prevented the PM_2.5_-induced increase in systemic TMAO and was associated with improved Y-maze performance and with rescue of Ki67+ cell loss in females. For DCX, PM_2.5_ treatment produced an overall effect, and sex-stratified comparisons suggest higher DCX+ counts in PM_2.5_-exposed FMO3*^cre/-^* mice in both sexes; however, because the sex × treatment interaction was not significant, these within-sex differences treated as exploratory rather than definitive evidence of sex-specific protection.

PERK, ATF4, and eIF2α levels were measured to assess ER stress signaling after PM_2.5_ exposure. Two□way ANOVA revealed no significant main effects or interactions for PERK, and ATF4 and eIF2α were similarly unchanged across groups (Fig. S6d–f). Although exploratory post hoc comparisons indicated lower PERK in PM_2.5_+FMO3*^cre/-^*versus PM_2.5_+FMO3*^flox/flox^* females, this difference was not supported by a significant ANOVA term. Overall, FMO3 deletion had minimal impact on ER stress signaling.

### PM_2.5_-induced hippocampal ER stress to induce cognitive and synaptic deficits

Chronic PM_2.5_ exposure disrupted hippocampal redox balance, as indicated by increased SOD activity and elevated MDA levels, consistent with oxidative membrane damage, whereas GSH-Px activity was unchanged (Fig. S6a). We therefore examined activation of the PERK branch of the unfolded protein response in the hippocampus. p-PERK, PERK, and BiP each showed significant main effects of treatment, with no significant effect of sex (Fig. 6b–d). ATF4 showed a trend toward a treatment effect, whereas eIF2α was unchanged (Fig. 6e, f). These data indicate that chronic PM_2.5_ exposure is associated with elevated oxidative stress and selective engagement of PERK-related ER stress signaling in the hippocampus.

**Fig. 6.**
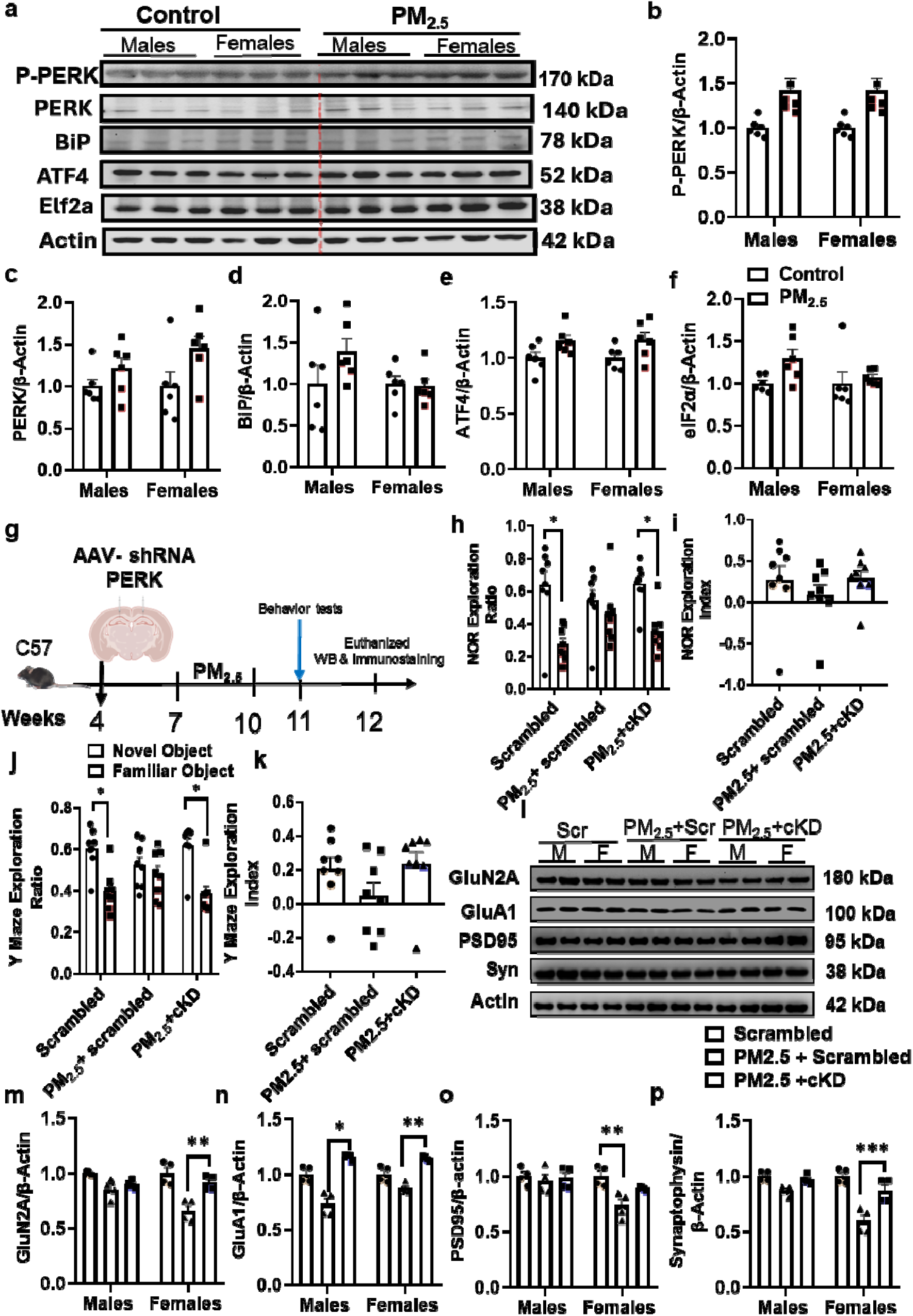
Hippocampal PERK inhibition attenuates behavioral impairment induced by chronic PM2.5 exposure. a-f. PM_2.5_ exposure activates PERK-associated ER stress signaling in the hippocampus. **a** Representative western blots. Quantification revealed a significant main effect of treatment on (**b**) p-PERK [F(1,5)=7.017, P=0.0455], (**c**) PERK [F(1,5)=16.66, P=0.0095], and (**d**) BiP [F(1,5)=6.890, P=0.0468], with no significant effects of sex or sex × treatment interaction. **e** ATF4 showed a trend toward a treatment effect [F(1,5)=6.002, P=0.0580], whereas (**f**) eIF2α was not significantly altered. n=6/group. **g** Experimental design and timeline. **h, i** Novel object recognition test: **h** Control scrambled mice, and PM_2.5_+cKD mice showed significant preference for the novel object, whereas PM_2.5_+scrambled mice did not (scrambled *P=0.0087, PM_2.5_+cKD P=0.018; paired t-tests). **i** One-way ANOVA showed that the discrimination index did not differ significantly among scrambled, PM_2.5_+scrambled, and PM_2.5_+cKD groups [F(2,21)=0.679, P=0.518]. **j-k** Y-maze performance. **j** In the arm exploration ratio, scrambled and PM_2.5_+cKD mice showed significant preference for the novel arm (paired t-tests; P=0.022 and P=0.018, respectively), whereas PM_2.5_+scrambled mice did not (P=0.605). **k** One-way ANOVA showed that the Y-maze exploration index did not differ significantly among scrambled, PM_2.5_+scrambled, and PM_2.5_+cKD groups [F(2,21)=1.854, P=0.181]. n=8/group. **l-p** Synaptic protein expression in the hippocampus. **l** Representative western blot of synaptic proteins. **m** GluN2A showed a significant main effect of treatment [F(1.058, 3.173)=18.19, P=0.0208], with no significant effects of sex or sex × treatment interaction. Post hoc analysis revealed a significant difference between PM_2.5_+cKD and PM_2.5_+scrambled females (*P=0.0014). **n** GluA1 also showed a significant main effect of treatment [F(1.012, 3.036)=69.75, *P=0.0034], with no significant effects of sex or sex × treatment interaction. Tukey’s multiple comparisons test showed significant differences between scrambled and PM_2.5_+scrambled males (P=0.0476), PM_2.5_+cKD and PM_2.5_+scrambled males (*P=0.0052), and PM_2.5_+cKD and PM_2.5_+scrambled females (*P=0.0020). **o** PSD95 showed significant effects of treatment [F(1.944, 5.832)=6.060, P=0.0382] and sex × treatment interaction [F(1.670, 5.009)=7.178, P=0.0362], and post hoc analysis identified a significant difference between scrambled and PM_2.5_+scrambled females (*P=0.0040). **p** SYN showed significant effects of sex [F(1,3)=16.50, P=0.0269], treatment [F(1.039, 3.118)=16.77, P=0.0243], and sex × treatment interaction [F(1.043, 3.129)=12.93, P=0.0342]. Post hoc comparisons revealed significant differences between scrambled and PM_2.5_+scrambled males (P=0.0268), scrambled and PM_2.5_+scrambled females (P=0.0393), and PM_2.5_+cKD and PM_2.5_+scrambled females (***P<0.0001). n=4 mice/group. All data are expressed as mean ± SEM.

To test the functional contribution of this pathway, we used a conditional PERK knockdown (cKD) in the hippocampus. In both the NOR and Y-maze tasks, PM_2.5_-exposed scrambled controls lost novelty preference, whereas PM_2.5_-exposed PERK cKD mice retained it (Fig. 6h, j). However, PERK knockdown partially preserved novelty-preference patterns, although group differences in exploration indices were not significant (Fig. 6i, k). We next examined hippocampal synaptic proteins. GluN2A and GluA1 showed significant main effects of treatment, with no significant sex or interaction effects (Fig. 6m, n). Although most GluN2A pairwise comparisons were not significant, PM_2.5_-exposed female cKD mice had higher GluN2A than PM_2.5_-exposed scrambled controls. For GluA1, PM_2.5_-exposed scrambled mice showed reduced levels, which were attenuated by PERK knockdown, particularly in males but also in females. PSD-95 and synaptophysin showed significant treatment effects and significant sex × treatment interactions (Fig. 6o, p). In females, PM_2.5_ reduced PSD-95, and PM_2.5_-exposed scrambled mice of both sexes showed lower synaptophysin than controls; notably, female cKD mice showed higher synaptophysin than PM_2.5_-exposed scrambled females. Together, these findings support a role for PERK signaling in PM_2.5_-induced hippocampal dysfunction and indicate that PERK knockdown partially preserves memory-related behavior and synaptic protein expression.

The effect on Ki67^+^ proliferating cells appears to be more evident in females, and cKD effectively rescued this early proliferative deficit in female mice. In contrast, although PM_2.5_ reduced DCX+ immature neurons in both sexes, cKD was not sufficient to significantly reverse this later-stage neurogenesis marker, suggesting that its protective effect may be stronger at the level of cell proliferation than at the stage of immature neuron survival or differentiation (Fig. S6e & f).

### Dietary reduction of ER stress ameliorates PM_2.5_-induced cognitive and synaptic impairment

To determine whether reducing ER stress mitigates the neurobehavioral effects of chronic PM_2.5_ exposure, we used a resveratrol (RSV) enriched diet. In the NOR task, control mice showed the expected preference for the novel object, whereas PM_2.5_-exposed mice lost this preference and instead favored the familiar object, indicating impaired working memory (Fig. 7a). RSV treatment restored novelty preference, indicating recovery of recognition memory in PM_2.5_-exposed mice. One-way ANOVA revealed significant group differences in the NOR exploration index (Fig. 7b); post hoc analysis showed that PM_2.5_ significantly reduced the index compared to controls, whereas RSV significantly increased it compared to PM_2.5_ alone. In the Y-maze, control mice preferentially explored the novel arm, while PM_2.5_ exposure shifted exploration toward the familiar arm (Fig. 7c). RSV reversed this effect, increasing time spent in the novel arm. One-way ANOVA confirmed significant group differences in the Y-maze exploration index, with PM_2.5_-exposed mice performing worse than controls (Fig. 7d). Together, these findings indicate that RSV attenuates PM_2.5_-induced deficits in both recognition memory and spatial novelty preference.

**Fig. 7.**
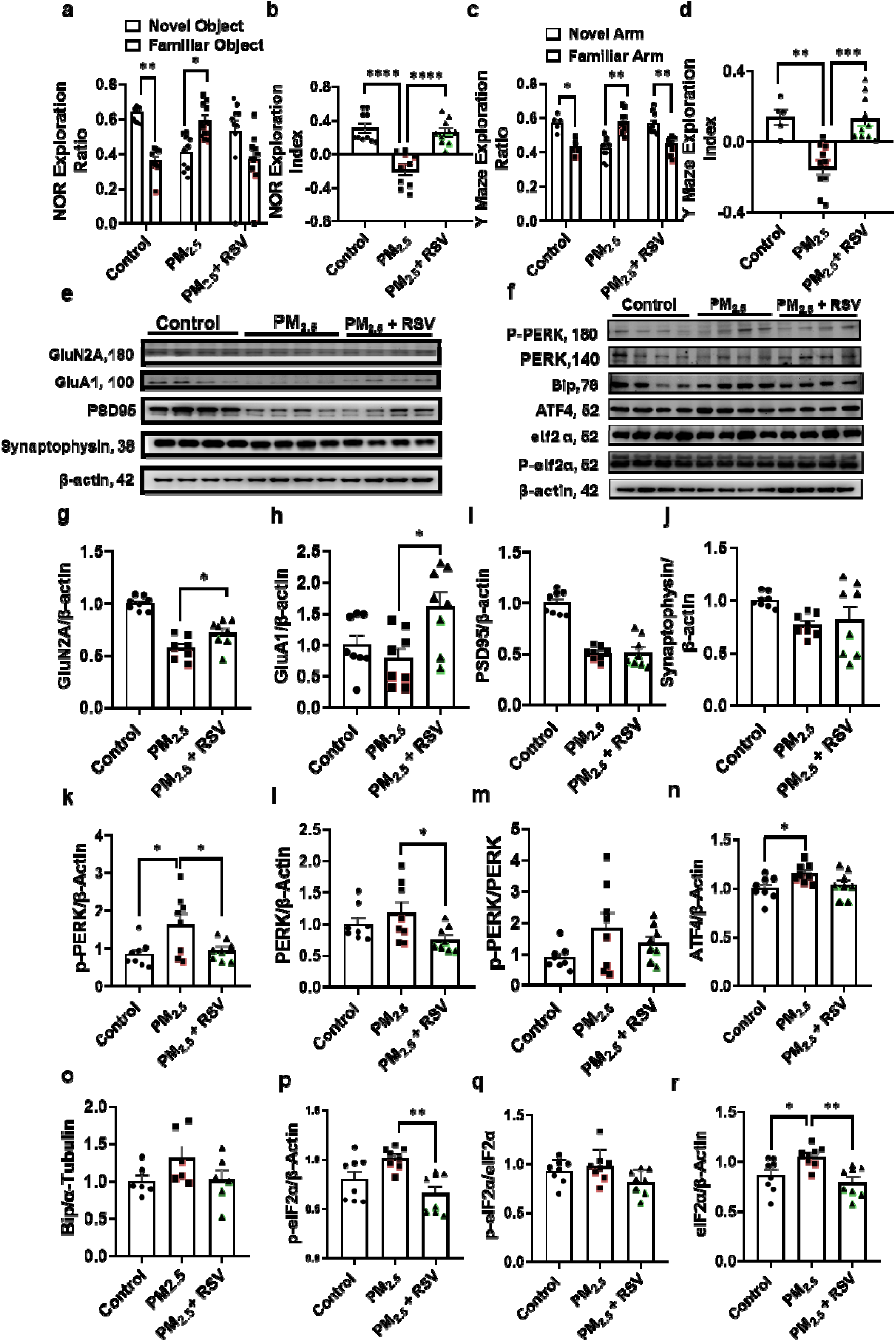
Reducing ER stress with dietary resveratrol attenuates chronic PM_2.5_-induced memory and hippocampal deficits. a-b. Novel object recognition test. In the object exploration ratio, control mice showed a significant preference for the novel object (**P=0.0020), whereas PM_2.5_ mice preferred the familiar object (*P=0.0185). **a** RSV treatment did not restore a significant preference for the novel object (P=0.1309). **b** One-way ANOVA showed significant differences in the discrimination index among groups (F=23.78, P<0.0001); PM_2.5_ significantly reduced the index compared with control (****P<0.0001), and this deficit was significantly improved by RSV treatment (***P<0.0001 vs. PM_2.5_). **c, d** Y-maze task. **c** In the arm exploration ratio, both control and RSV mice spent significantly more time in the novel arm (*P=0.0312 and **P=0.0034, respectively), whereas PM_2.5_ treatment significantly increased exploration of the familiar arm (**P=0.0038). **d** One-way ANOVA revealed significant group differences in the Y-maze exploration index (F=15.24, P<0.0001), with PM_2.5_ mice showing reduced performance relative to control (**P=0.0011) and RSV significantly improving this deficit (***P=0.0002 vs. PM_2.5_). n=10/group. **e, g-j** Hippocampal synaptoneurosome analysis. Representative western blots for synaptic markers are shown in (**e**). **g** GluN2A was partially restored by RSV treatment (*P=0.0346), and GluA1 was significantly increased compared with the PM_2.5_ group (**h**; F=5.568, P=0.0115; post hoc P=0.0111). No significant changes were detected in PSD95 (**i**) or Synaptophysin (**j**; F=2.662, P=0.0933). **f**, **k-r** ER stress-related proteins in the hippocampus. Representative western blots of PERK pathway proteins are shown in (**f**). PM_2.5_ significantly increased p-PERK (**k**; P=0.0171), ATF4 (**n**; P=0.0454), and eIF2α (**r**; P=0.0362). RSV treatment significantly reduced p-PERK (**k**; P=0.0390) and eIF2α (**r**; *P=0.0046) and increased total PERK (**l**; P=0.0499) and p-eIF2α (**p**; *P=0.0024) compared with the PM_2.5_ group. No significant differences were observed in the p-PERK/PERK ratio (**m**), BiP (**o**), or the p-eIF2α/eIF2α ratio (**q**). n=8/group. All data are expressed as mean ± SEM.

We next examined whether these behavioral improvements were accompanied by restoration of synaptic protein expression. RSV exerted selective effects on glutamatergic synaptic markers (Fig. 7g–j). GluN2A expression was partially restored in RSV-treated mice compared to the PM_2.5_ group (Fig. 7g), indicating partial recovery of NMDA receptor-associated signaling. In contrast, GluA1 was significantly increased by RSV (Fig. 7h), suggesting more robust rescue of AMPA receptor-related signaling. However, RSV did not restore PSD95 expression (Fig. 7i), and synaptophysin levels remained unchanged across groups (Fig. 7j). Thus, RSV appears to selectively preserve glutamatergic receptor subunits rather than normalizing all synaptic components altered by PM_2.5_ exposure.

Western blotting analysis revealed that PM_2.5_ altered the PERK downstream cascade, which was partially reversed by RSV (Fig. 7k–r). Specifically, RSV reduced both p-PERK and total PERK compared to PM_2.5_ exposure alone (Fig. 7k, l). However, the p-PERK/PERK ratio was unchanged among groups (Fig. 7m), suggesting that RSV primarily reduced PERK abundance rather than altering its relative phosphorylation state. RSV significantly reduced downstream eIF2α levels (Fig. 7n), whereas ATF4 remained unchanged (Fig. 7r), indicating only partial suppression of the canonical PERK pathway. Although p-eIF2α did not differ significantly between control and PM_2.5_ groups, RSV significantly reduced p-eIF2α relative to PM_2.5_ alone (Fig. 7p), while the p-eIF2α/eIF2α ratio remained unchanged (Fig. 7q). Collectively, these findings suggest that RSV does not uniformly normalize the entire PERK pathway, but instead selectively dampens key components of the ER stress response associated with PM_2.5_ exposure. Taken together, these data indicate that RSV is effective in protecting against PM_2.5_-induced hippocampal dysfunction, as reflected by improved memory performance, selective preservation of glutamatergic synaptic proteins, and attenuation of PERK-associated ER stress signaling.

## Discussion

This study establishes a novel mechanistic framework for PM_2.5_-induced neurotoxicity, demonstrating that chronic exposure impairs cognitive function via a gut–liver–brain axis. While the link between air pollution and neurodegeneration is well recognized, the systemic mediators linking pulmonary inhalation to hippocampal dysfunction remain poorly defined. Our findings identify the microbial metabolite TMAO as the central mediator of this process. Specifically, we show that PM_2.5_ exposure disrupts the gut microbial landscape, favoring taxa responsible for TMA synthesis. This microbial shift drives a systemic increase in TMAO production in the liver, which subsequently accumulates in the hippocampus, triggering PERK-mediated ER stress. This cascade ultimately culminates in impaired adult hippocampal neurogenesis, impaired synaptic plasticity, and memory deficits. These results not only broaden our understanding of the systemic nature of air pollution-induced neurodegeneration but also identify microbial metabolites as viable therapeutic targets for mitigating the neurological consequences of environmental pollutants.

Chronic PM_2.5_ exposure significantly impaired cognitive function without altering basal locomotor activity, suggesting that these deficits reflect specific neurological impairment. Specifically, PM_2.5_-exposed mice exhibited a loss of novelty preference in both the novel object recognition and Y-maze tasks, consistent with established reports that fine particulate matter disrupts memory-related behaviors ^6,62–65^. Our previous studies have indicated that cognitive impairment is linked to hippocampal dysfunction, including reduced adult neurogenesis, dendritic branching, spine density, and synaptic plasticity ^66–68^. Similarly, we found that PM_2.5_-induced behavioral deficits were underpinned by reduced hippocampal synaptic plasticity, attenuated LTP, and diminished expression of key synaptic proteins, including GluA1, PSD95, and synaptophysin. These findings corroborate earlier studies linking PM_2.5_ to structural and functional hippocampal degradation ^6,69^, while extending the literature by demonstrating that behavioral decline is tightly coupled with electrophysiological and molecular evidence of compromised hippocampal structural and functional plasticity.

Another key finding of this study is that PM_2.5_ disrupts the gut microbiome in a manner that favors TMA production, subsequently elevating TMAO levels in both the systemic circulation and the hippocampus. Our metagenomic analysis revealed broad shifts in microbial community structure and a specific enrichment of genes involved in choline utilization (cutC and cutD). While previous research has shown that PM_2.5_ perturbs gastrointestinal homeostasis, promoting inflammation and dysbiosis through dysregulated autophagy or altered β-diversity ^70^. These findings align with a growing body of evidence indicating that particulate air pollution disrupts gastrointestinal homeostasis and microbial ecology. Previous studies have demonstrated that ambient PM_2.5_ compromises colonic health by promoting inflammation, abnormal proliferation, and dysbiosis, potentially via dysregulated autophagy ^71^. Similarly, research utilizing diesel PM_2.5_ intratracheal instillation reported significant alterations in microbial diversity and community structure, characterized by distinct β-diversity shifts and changes in key taxa associated with inflammatory and metabolic dysregulation ^72^. Inhalation models have yielded comparable results, showing that concentrated ambient particles remodel the microbial landscape across the gastrointestinal tract ^73^. Our data corroborate these reports of PM_2.5_-induced microbial disruption, extending current knowledge by identifying a specific functional consequence. The enrichment of microbial genes driving choline-to-TMA metabolism and a concomitant rise in systemic TMAO levels. This frames dysbiosis as a mechanistic, gut-derived pathway linking PM_2.5_ exposure to hippocampal dysfunction.

Our intervention studies provide compelling evidence that TMAO is a functional driver of PM_2.5_-induced cognitive impairment. Pharmacological inhibition of microbial TMA production via DMB attenuated TMAO accumulation and ameliorated deficits in memory, neurogenesis, and synaptic plasticity. Complementary results were obtained through genetic disruption of FMO3, the host enzyme responsible for converting TMA to TMAO, thereby preventing the systemic TMAO surge and preserving hippocampal function. The convergence of these two distinct strategies is critical; by targeting the same metabolic pathway at different stages (microbial production vs. host hepatic conversion), we strengthen the conclusion that TMAO is a mechanistic mediator of neurotoxicity rather than a mere correlation of exposure. This aligns with clinical evidence associating elevated TMAO with cognitive decline in older adults ^74^ and experimental data showing that TMAO exacerbates neurodegeneration via oxidative stress and SIRT1 suppression ^75,76^. While the breadth of the rescue across behavioral and electrophysiological measures was notable, the protection remained partial, suggesting that while the TMAO pathway is a primary mediator, it operates alongside other contributing factors in PM_2.5_-induced brain injury.

Furthermore, we identify hippocampal ER stress, specifically the PERK signaling pathway, as a critical downstream effector of the PM_2.5_–TMAO axis. Our results demonstrate that PM_2.5_ exposure upregulates key components of the PERK pathway, including BiP, PERK, and p-PERK, and elevates oxidative stress markers. This suggests that pollutant exposure triggers a stress response linked to redox imbalance. Crucially, our data indicates that this ER stress response is driven by a rise in TMAO. This is also supported by biochemical evidence that TMAO can directly bind to and activate PERK ^77^, as well as omics-based studies identifying TMAO as a primary driver of ER stress during glutathione depletion ^78^. Furthermore, our findings align with established models of liver and cardiac injury, where TMAO-induced PERK activation orchestrates tissue dysfunction. Evidence from other organ systems supports the same mechanism. In liver injury models, TMAO activates PERK signaling, and inhibition of TMAO production attenuates ER stress and tissue injury ^79^ ^80^. By demonstrating that hippocampal-specific PERK knockdown provides neuroprotection, we establish that TMAO-driven ER stress is a major mechanistic link between air pollution and the loss of synaptic integrity.

The influence of biological sex on the response to PM_2.5_ is a critical aspect of our findings. While PM_2.5_ impaired memory in both sexes, female mice exhibited more pronounced reductions in hippocampal cell proliferation and immature neurons, along with more significant shifts in microbial α-diversity. Furthermore, females showed a more robust rescue of several endpoints following DMB treatment and clearer rescuing effects of Ki67+ cells after FMO3 disruption. These patterns align with emerging clinical and epidemiological evidence suggesting that females may be particularly susceptible to the adverse effects of air pollution. For instance, in a large cohort of middle-aged and elderly individuals, the association between PM_2.5_ exposure and cognitive decline was notably more pronounced in women ^81^. Similarly, chronic PM_2.5_ has been linked to Alzheimer’s disease–like cognitive impairment preferentially in females ^82^. Our findings extend this literature by suggesting that this heightened female vulnerability is not only behavioral but is also reflected in physiological domains tied to hippocampal integrity and gut–brain signaling. The disproportionate response to TMAO-lowering interventions in females suggests that the TMA/TMAO pathway may be a primary driver of sex-dependent neurotoxicity, potentially reflecting underlying differences in microbial ecology, hepatic metabolic handling of TMA, or baseline hippocampal redox regulation.

Collectively, these findings provide mechanistic insight into air pollution neurotoxicity. Our data support the role of the gut–liver–brain axis involved in the detrimental effects of PM_2.5_ on brain health, by which inhaled PM_2.5_ alters the gut microbiome, enriches for TMA-producing taxa, increases systemic TMAO, and ultimately activates a PERK-dependent stress that consequently impairs hippocampal plasticity. Our findings have significant translational implications, offering potential interventions. Our results with DMB, FMO3 modulation, and RSV treatment demonstrate the potential of targeting microbial metabolism to preserve cognitive health. While these interventions may not fully neutralize the multifaceted effects of air pollution, our study identifies microbial TMAO production and hippocampal PERK signaling as high-priority targets to reduce neurological risk. More importantly, our findings integrate environmental health with the growing field of microbial and brain health, providing a clear example of how gut-derived metabolites can shape brain vulnerability to environmental stressors.

This study has several limitations that warrant consideration. First, we employed intratracheal instillation rather than whole-body inhalation exposure. While instillation allows for precise dose control and ensures that the observed effects are triggered by the pulmonary presence of particles, it may not fully replicate the dynamic deposition patterns of real-world aerosols. However, given that the downstream shifts in the microbiome, systemic metabolites, and hippocampal pathology were internally consistent across all our intervention models, it is unlikely that the mode of delivery alters the fundamental gut–liver–brain pathway identified here. Second, while our mechanistic analysis focused on PERK signaling, PM_2.5_ exposure almost certainly recruits parallel pathways, including neuroinflammation, vascular dysfunction, and other branches of the unfolded protein response. The multifaceted nature of PM_2.5_ neurotoxicity likely explains why TMAO suppression and PERK knockdown provided significant, but not absolute, rescue of the cognitive phenotype. Future studies utilizing fecal microbiota transplantation from PM_2.5_-exposed donors into germ-free or antibiotic-depleted recipients will be essential. Such experiments would directly test whether the altered microbiome is sufficient to transmit the neurobehavioral phenotype, thereby further refining the causal inference at the gut–brain interface.

In conclusion, this study establishes a novel mechanistic framework for PM_2.5_-induced cognitive impairment, mediated by a gut–liver–brain axis. Our findings demonstrate that inhaled fine particulates modulate gut microbial metabolism, favoring the synthesis of the metabolite TMAO, which subsequently activates the hippocampal PERK-dependent ER stress pathway, impairing synaptic plasticity and memory. The efficacy of targeting microbial TMA production, hepatic FMO3 conversion, and hippocampal PERK signaling in rescuing these deficits, complemented by the protective effects of dietary RSV, provides robust evidence for the causality of this systemic metabolic relay. Furthermore, the identification of a more pronounced vulnerability in females underscores the necessity of sex-specific considerations in environmental neurotoxicology. By shifting the paradigm of air pollution neurotoxicity from a localized inflammatory response toward a systemic, multi-organ metabolic framework, this work reveals mechanistic insights that may be targeted to preserve cognitive resilience in the face of escalating global air pollution.

## Acknowledgement

We would like to thank the research support from the Centralized Animal Facility (CAF) and the University Research Facility in Behavioral and Systems Neuroscience (UBSN).

