## Supplementary figures for "Microbiota-derived TMAO links PM_2.5_ exposure to cognitive impairment: A gut–liver–brain axis"

Hussain et al

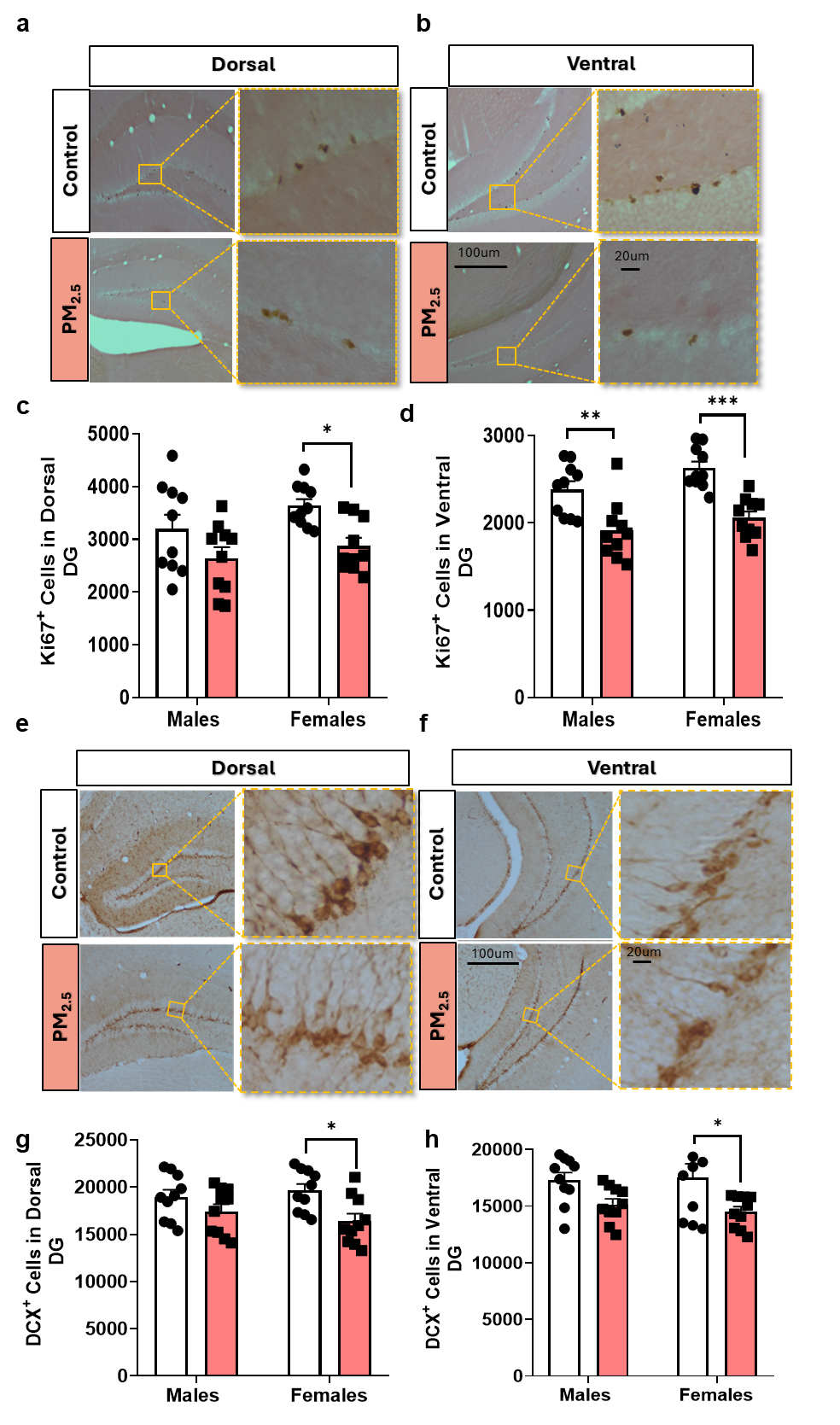

**Fig. S1. Chronic PM_2.5_ exposure suppresses hippocampal neurogenesis in the dentate gyrus. a, b** Representative images of Ki67-immunolabeled proliferating cells in the dorsal (**a**) and ventral dentate gyrus (DG) (**b**) of control and PM_2.5_-exposed mice. **c, d** Quantification of Ki67-positive cells in the dorsal DG (**c**) and ventral DG (**d**) showed that chronic PM_2.5_ exposure significantly reduced proliferative activity. In the dorsal DG (**c**), two-way ANOVA revealed significant main effects of sex [F(1, 9) = 6.224, P = 0.0342] and treatment [F(1, 9) = 43.27, P = 0.0001], with no significant sex × treatment interaction [F(1, 9) = 0.4328, P = 0.5271]; Tukey’s multiple-comparisons test showed a significant reduction in female mice (P = 0.0067), whereas the reduction in male mice did not reach significance (P = 0.0907). In the ventral DG (**d**), two-way ANOVA showed significant main effects of sex [F(1, 9) = 7.250, P = 0.0247] and treatment [F(1, 9) = 45.93, P < 0.0001], with no significant interaction [F(1, 9) = 0.9707, P = 0.3503]; Tukey’s multiple-comparisons test showed significant reductions in both male (P = 0.0015) and female (P = 0.0014) mice. **e, f** Representative images of DCX-immunostained immature neurons in the dorsal (**e**) and ventral DG (**f**) of control and PM_2.5_-exposed mice. **g, h** Quantification of DCX-positive cells in the dorsal DG (**g**) and ventral DG (**h**) demonstrated a significant overall effect of PM_2.5_ exposure on immature neuron abundance. In the dorsal DG (**g**), two-way ANOVA showed a significant main effect of treatment [F(1, 9) = 9.951, P = 0.0117], with no significant effects of sex [F(1, 9) = 0.1240, P = 0.7328] or sex × treatment interaction [F(1, 9) = 1.874, P = 0.2042]. In the ventral DG (**h**), two-way ANOVA also showed a significant main effect of treatment [F(1, 9) = 9.984, P = 0.0116], with no significant effects of sex [F(1, 9) = 0.1473, P = 0.7101] or interaction [F(1, 9) = 0.6687, P = 0.4346]; Tukey’s multiple-comparisons test showed a significant reduction in male mice (P = 0.0223), whereas the decrease in female mice did not reach statistical significance (P = 0.1410). Scale bars: 100 μm for main images and 20 μm for insets. n=10/group. Data are presented as mean ± SEM with individual data points.

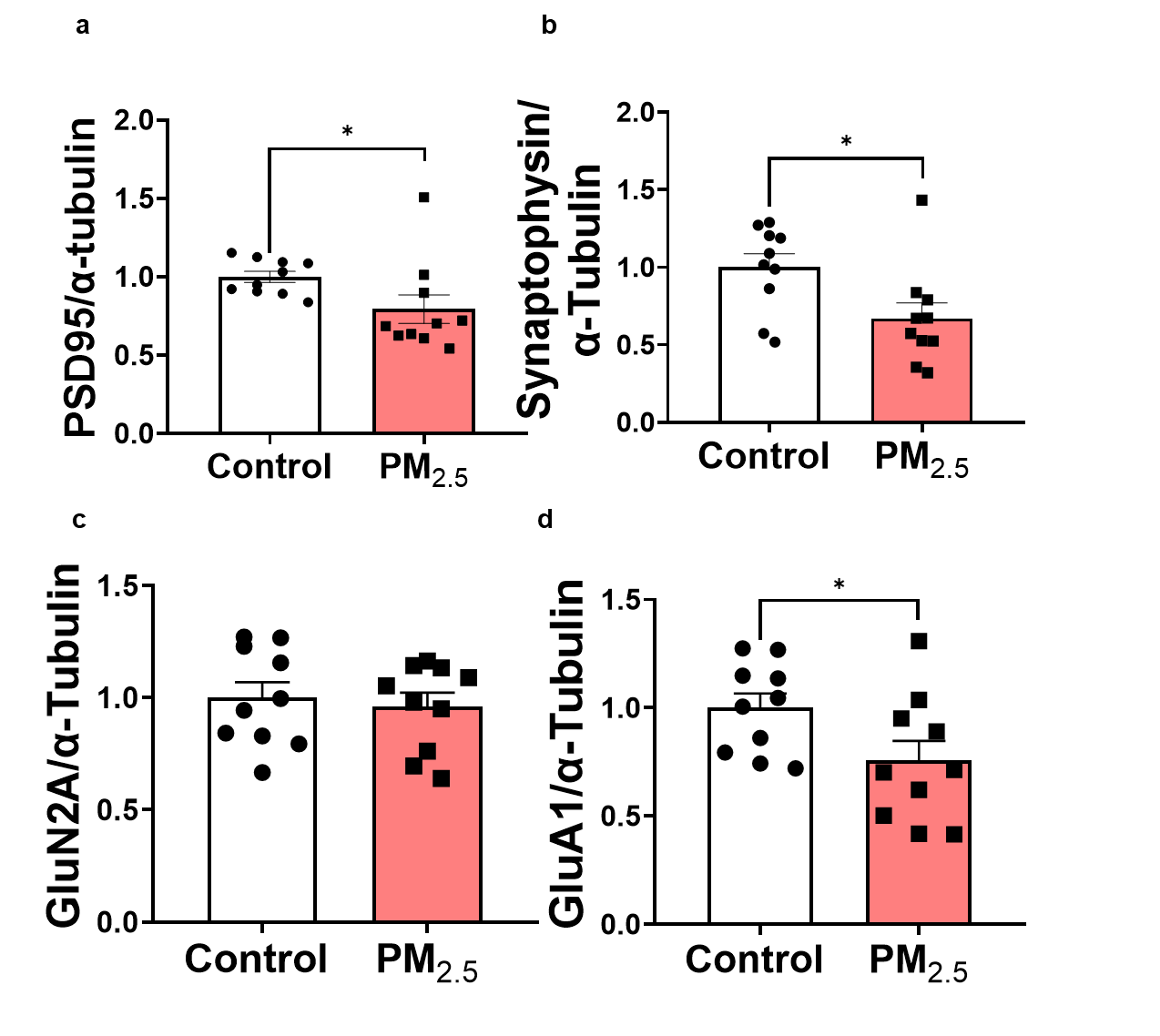

**Fig. S2. Chronic PM_2.5_ exposure reduces the expression of key synaptic proteins in the hippocampus.** **a-d** Quantification of synaptic protein levels revealed that chronic PM_2.5_ exposure significantly downregulated several synaptic plasticity markers. Specifically, PM2.5 exposure significantly reduced the expression of (**a**) PSD95 (P = 0.0115), (**b**) SYN (P = 0.0355), and (**d**) GluA1 (P = 0.0433). (**c**) In contrast, no significant difference was observed in GluN2A expression between the control and PM_2.5_ groups (P = 0.6305). n=10/group. Data are presented as mean ± SEM with individual data points.

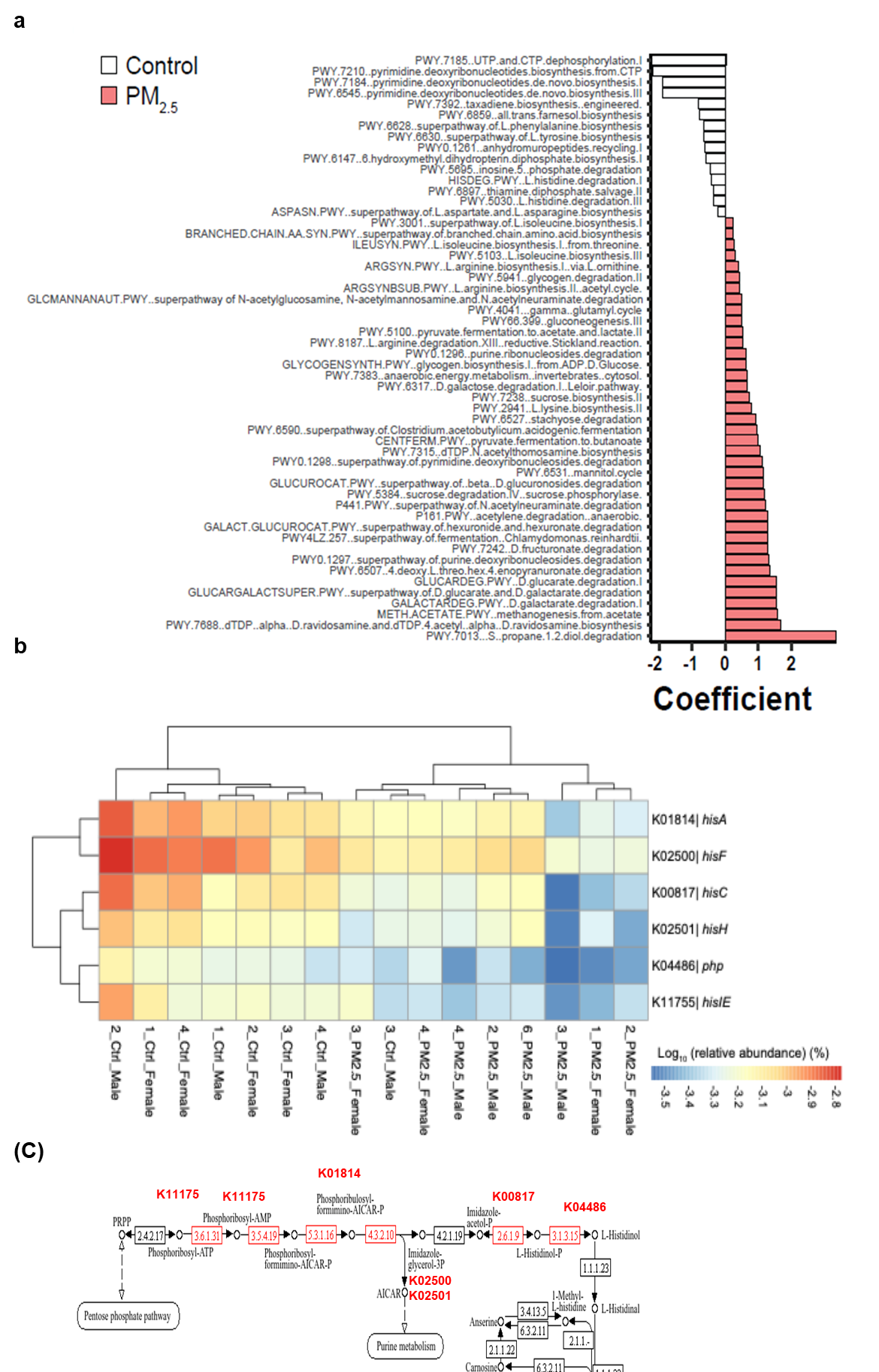

**Fig. S3. Chronic PM_2.5_ exposure reshapes microbial metabolic pathways and increases circulating TMAO.** **a** MaAsLin2 analysis identified significantly altered gut microbial metabolic pathways in PM_2.5_-exposed mice relative to control mice. Positive coefficients indicate pathways enriched after exposure, with prominent changes in amino acid and carbohydrate metabolism. **b** Heatmap showing differential abundance of genes involved in histidine and purine biosynthesis. **c** Hierarchical clustering of KEGG orthologs, including K01814 and K02500, revealed distinct pathway-associated gene profiles between groups, with red indicating higher abundance in the control group.

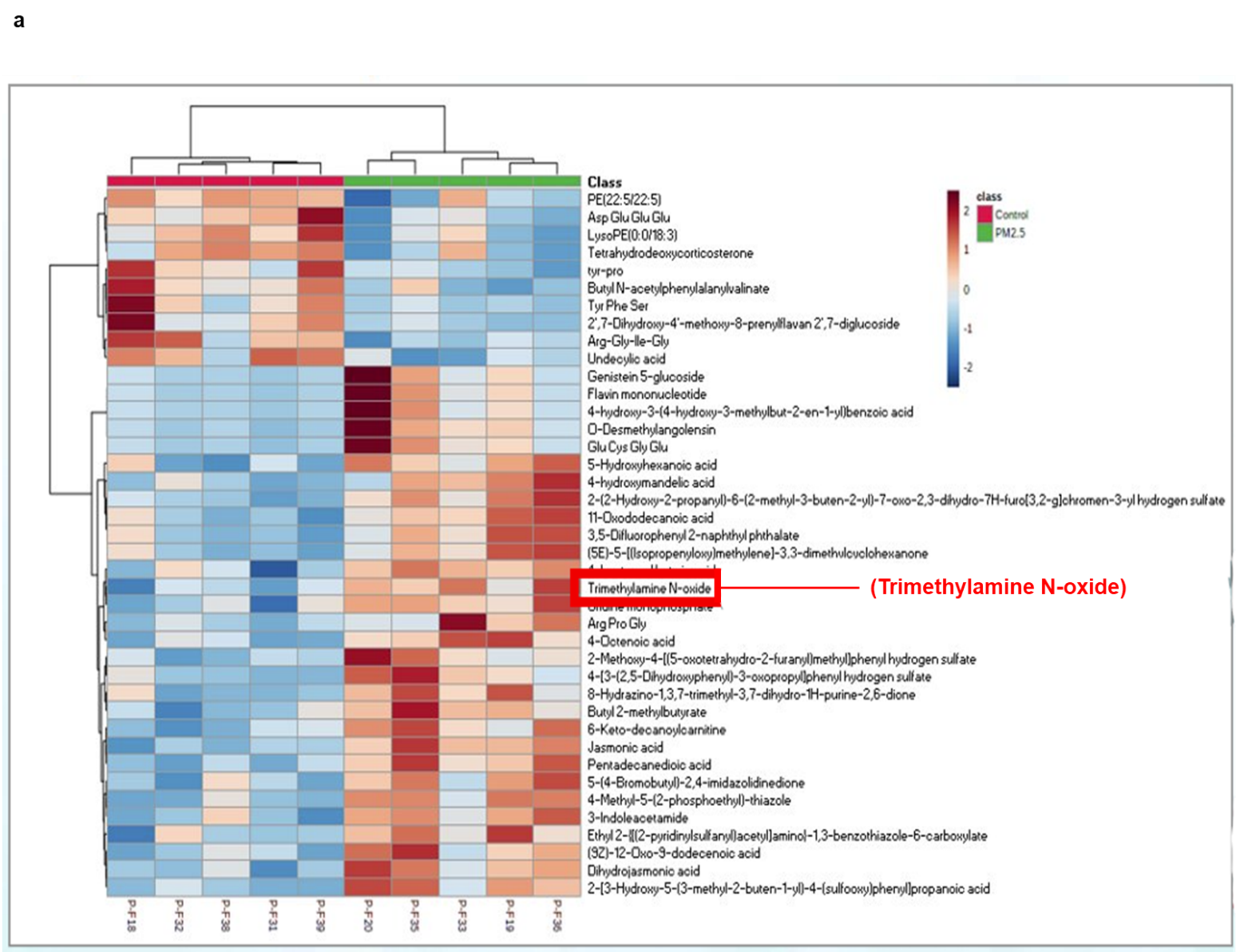

**Fig. S4. Heatmap of metabolite abundance in Control and PM_2.5_ groups.**
Hierarchical clustering heatmap showing relative levels of metabolites across individual samples from the Control and PM_2.5_ groups. Columns represent samples and rows represent metabolites. The top annotation bar indicates class membership (green, Control; red, PM_2.5_). Colors represent normalized metabolite abundance (z-score), with red indicating higher and blue indicating lower relative levels. The highlighted metabolite, trimethylamine N-oxide (TMAO), was differentially expressed between groups and may reflect PM_2.5_-associated alterations in microbial-host metabolic pathways.

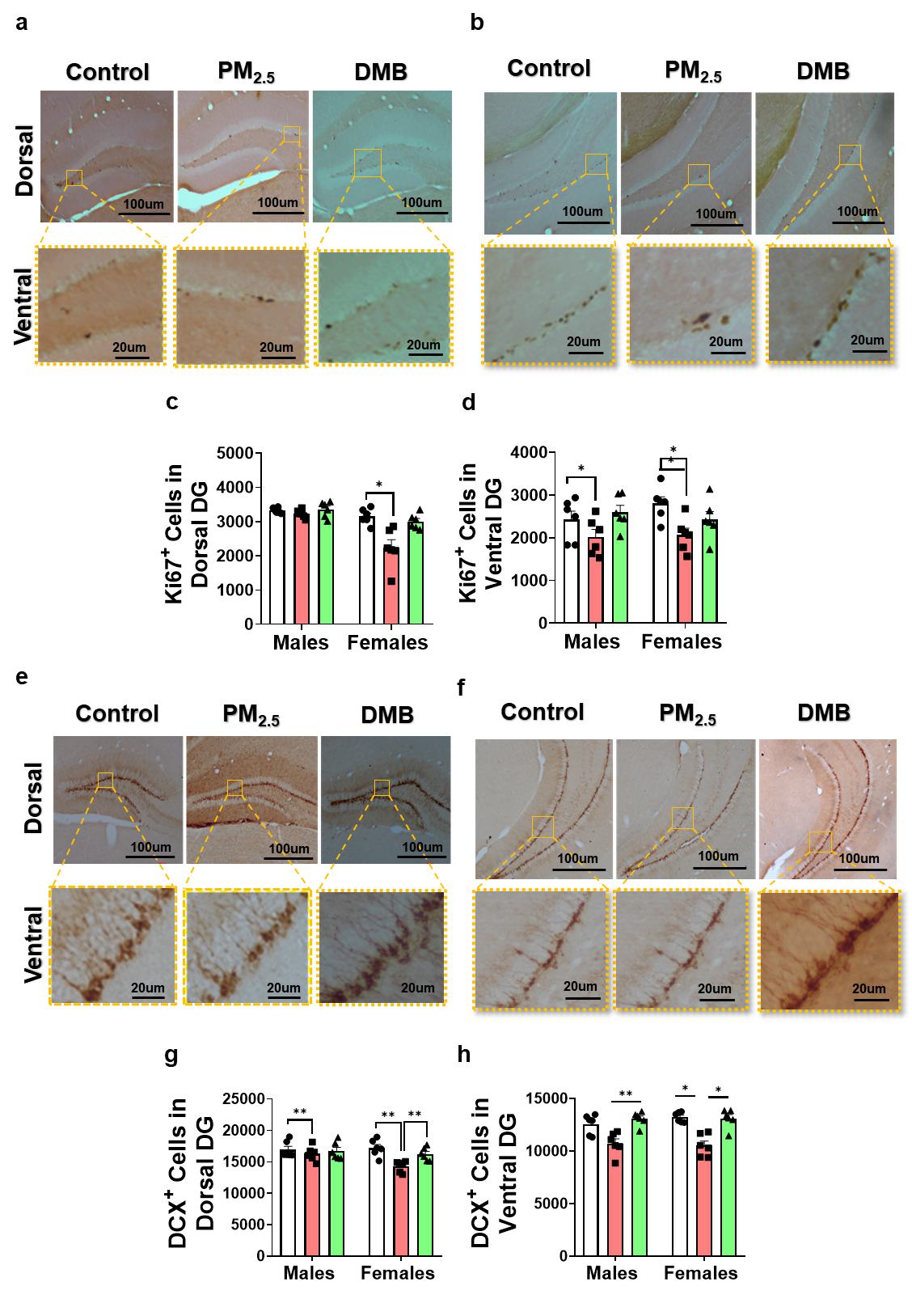

**Fig. S5. Inhibition of TMAO mitigates chronic PM_2.5_-induced suppression of hippocampal neurogenesis.** **a, b** Representative images of Ki67-immunostained proliferating cells in the dorsal (**a**) and ventral dentate gyrus (DG) (**b**). **c, d** Quantification of Ki67-positive cells in the dorsal DG (**c**) and ventral DG (**d**) showed that PM_2.5_ exposure reduced cell proliferation, whereas DMB treatment attenuated this effect. In the dorsal DG, two-way ANOVA revealed significant main effects of sex [F(1, 5) = 14.60, P = 0.0124] and treatment [F(1.211, 6.053) = 13.06, P = 0.0095], as well as a significant sex × treatment interaction [F(1.799, 8.995) = 9.470, P = 0.0069]; Tukey’s multiple-comparisons test showed a significant reduction in female mice (Control vs. PM_2.5_, P = 0.0172) and a significant rescue by DMB (PM_2.5_ vs. DMB, P = 0.0383), with no significant changes in male mice. In the ventral DG, two-way ANOVA showed a significant main effect of treatment [F(1.302, 6.512) = 16.94, P = 0.0040], with no significant effect of sex [F(1, 5) = 0.3672, P = 0.5710] or sex × treatment interaction [F(1.798, 8.992) = 3.841, P = 0.0654]; post hoc analysis showed significant PM_2.5_-induced reductions in both male (P = 0.0100) and female (P = 0.0144) mice. Post hoc testing indicated that DMB significantly increased proliferation relative to PM2.5 alone in males (P = 0.0433). **e, f** Representative images of DCX-immunostained immature neurons in the dorsal (**e**) and ventral DG (**f**). **g, h** Quantification of DCX-positive cells in the dorsal DG (**g**) and ventral DG (**h**) demonstrated that PM_2.5_ exposure reduced immature neuron abundance and that DMB partially reversed this effect. In the dorsal DG, two-way ANOVA showed a significant main effect of treatment [F(1.557, 7.785) = 32.99, P = 0.0002] and a significant sex × treatment interaction [F(1.732, 8.661) = 11.18, P = 0.0048], with no significant main effect of sex [F(1, 5) = 3.184, P = 0.1344]; Tukey’s multiple-comparisons test showed a significant reduction in female mice following PM_2.5_ exposure (P = 0.0009), which was significantly rescued by DMB (P = 0.0030), whereas no significant changes were observed in male mice. In the ventral DG, two-way ANOVA revealed a significant main effect of treatment [F(1.342, 6.712) = 18.56, P = 0.0028], with no significant effects of sex [F(1, 5) = 0.3139, P = 0.5995] or sex × treatment interaction [F(1.155, 5.774) = 2.216, P = 0.1910]; post hoc analysis showed a significant reduction in female mice after PM_2.5_ exposure (P = 0.0044), while DMB significantly increased DCX-positive cell numbers relative to the PM_2.5_ group in both male (P = 0.0029) and female (P = 0.0152) mice. Scale bars: 100 μm for main images and 20 μm for insets. n = 10 per group. Data are presented as mean ± SEM with individual data points.

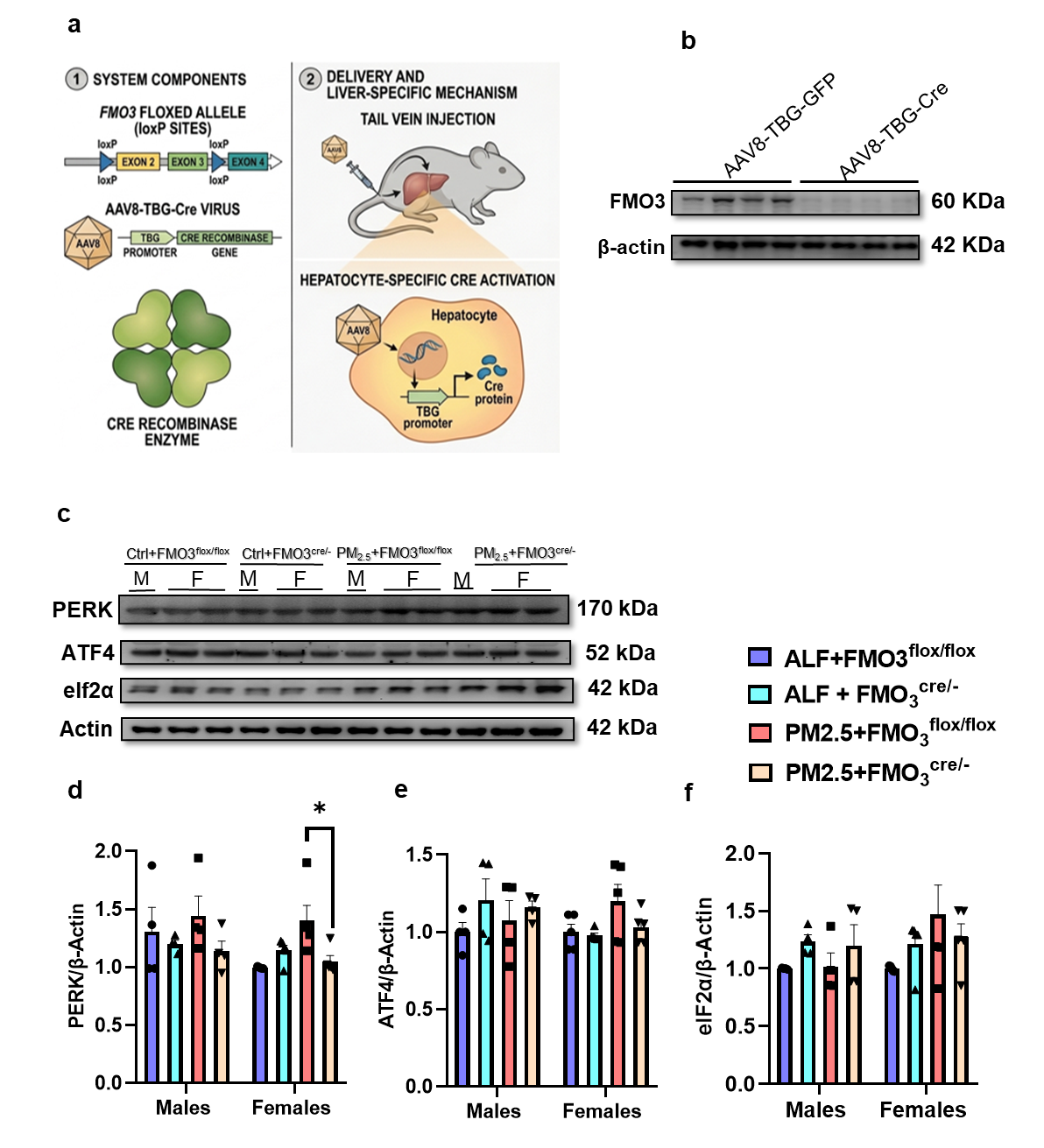

**Fig. S6. Hepatocyte-specific FMO3 deletion strategy and downstream PERK-related molecular analyses.** **a** Schematic illustration of the FMO3 floxed allele and the AAV8-TBG-Cre strategy used to achieve hepatocyte-specific deletion of FMO3. Cre recombinase, driven by the hepatocyte-specific TBG promoter, was delivered by tail-vein injection of AAV8-TBG-Cre, resulting in liver-specific deletion of FMO3 in hepatocytes. **b** Representative immunoblot showing validation of hepatic FMO3 reduction in the indicated experimental groups. **c** Representative immunoblots of ER stress marker proteins. **d**-**f** Quantification of the indicated ER stress-related proteins. **d** PERK levels showed no significant main effect of sex [F(1.000, 4.000)=2.210, P=0.2113], treatment [F(1.365, 5.459)=3.993, P=0.0925], or sex × treatment interaction [F(1.493, 3.982)=0.7387, P=0.4951]. Post hoc Tukey tests revealed that PERK levels were significantly lower in females in the PM2.5 + FMO3*^cre/-^* group compared with the PM2.5 + FMO3*^flox/flox^* group (*p=0.0477), whereas no significant pairwise differences were observed in males. **e** ATF4 levels showed no significant main effect of sex [F(1,28)=0.988, P=0.3290], group [F(3,28)=1.024, P=0.3976], or sex × group interaction [F(3,28)=1.728, P=0.1845]. Post hoc Tukey tests showed no significant differences among groups in either males or females. **f** eIF2α levels showed no significant main effect of sex [F(1.000, 4.000)=0.7119, P=0.4463], treatment [F(1.145, 4.581)=2.550, P=0.1781], or sex × treatment interaction [F(1.717, 4.579)=2.293, P=0.2033]. Tukey’s multiple comparisons test revealed no significant pairwise differences in either sex. n=4/group. Data are presented as mean ± SEM with individual data points shown.

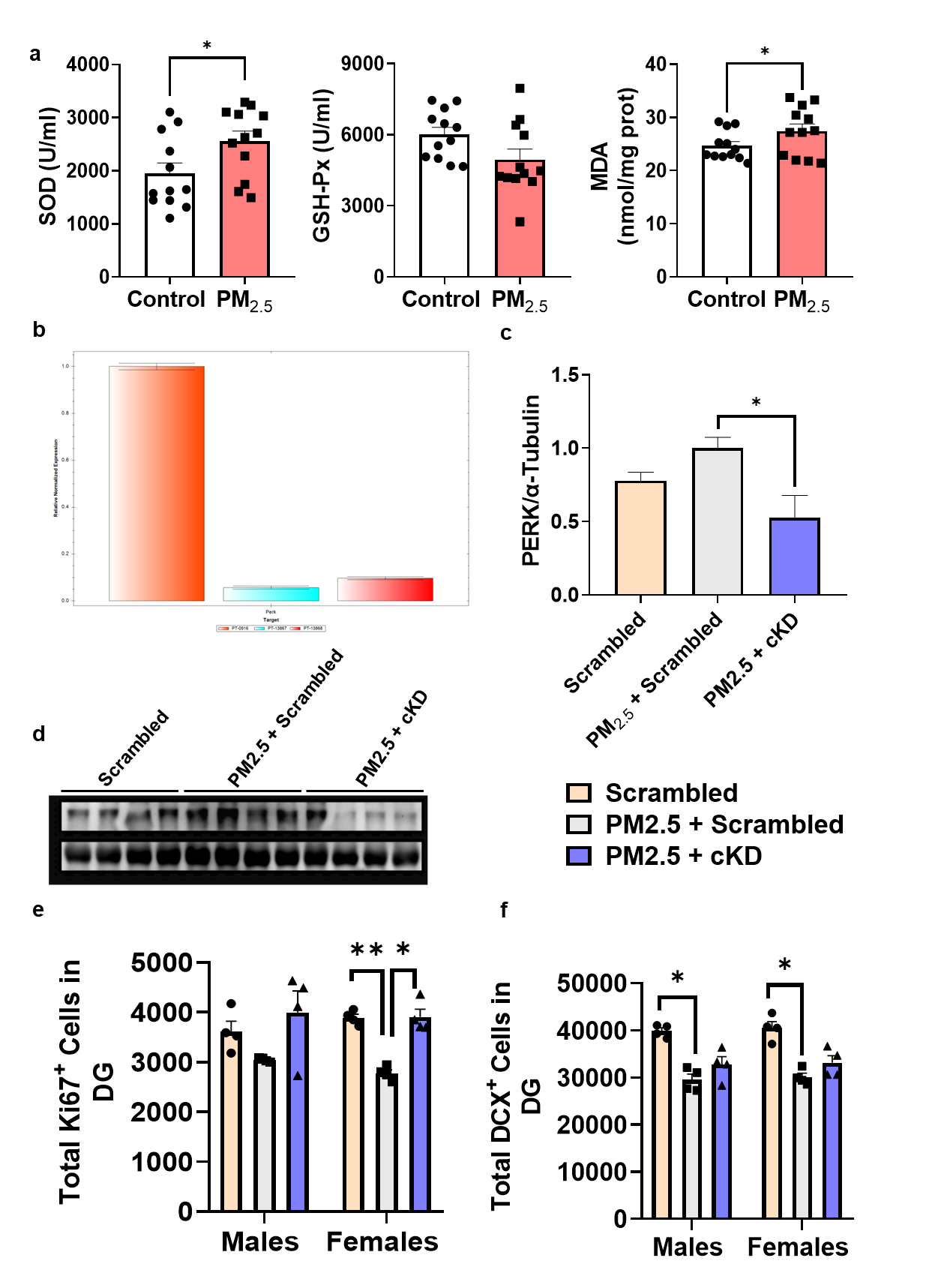

**Fig. S7. Chronic PM2.5 exposure induces oxidative stress, and PERK inhibition partially rescues dentate gyrus neurogenesis.** **a** Quantification of oxidative stress-related markers in the hippocampus, including SOD, GSH-Px, and MDA levels. Unpaired t test with Welch’s correction showed that SOD and MDA levels were significantly altered in the PM_2.5_ group compared with the control group, SOD (t = 2.249, df = 21.93, P = 0.0349), and MDA (t = 2.634, df = 19.40, P = 0.0162) whereas GSH-Px (t = 2.003, df = 19.47, P = 0.0593) were not significantly different. n=12/group. **b-d** Validation of AAV2/9-shPERK knockdown efficiency, including the experimental strategy and representative western blot analysis of PERK protein expression in the indicated groups. **e, f** Quantification of Ki67-positive proliferating cells and DCX-positive immature neurons in the dentate gyrus (DG). **e** Two-way ANOVA did not detect significant main effects of sex or treatment, nor a significant sex × treatment interaction. However, post hoc comparisons suggested a female-specific reduction in Ki67-positive cells in the PM2.5 + scrambled group relative to scrambled controls, and a corresponding increase in the PM2.5 + cKD group relative to PM2.5 + scrambled females. No significant pairwise differences were observed in males. Given the small sample size, these findings should be interpreted cautiously. **f** DCX-positive cell number. Two-way ANOVA revealed a significant main effect of treatment, with no significant main effect of sex and no significant sex × treatment interaction. Post hoc comparisons confirmed that PM2.5 + scrambled significantly reduced DCX-positive cell numbers relative to scrambled controls in both males and females, indicating that PM2.5 suppressed immature neuron abundance across sexes. However, no significant pairwise differences were detected between the PM2.5 + scrambled and PM2.5 + cKD groups in either sex, suggesting that PERK knockdown did not significantly rescue this endpoint under the conditions tested. Males: n=3/group; females: n=4/group. Data are presented as mean ± SEM with individual data points shown.

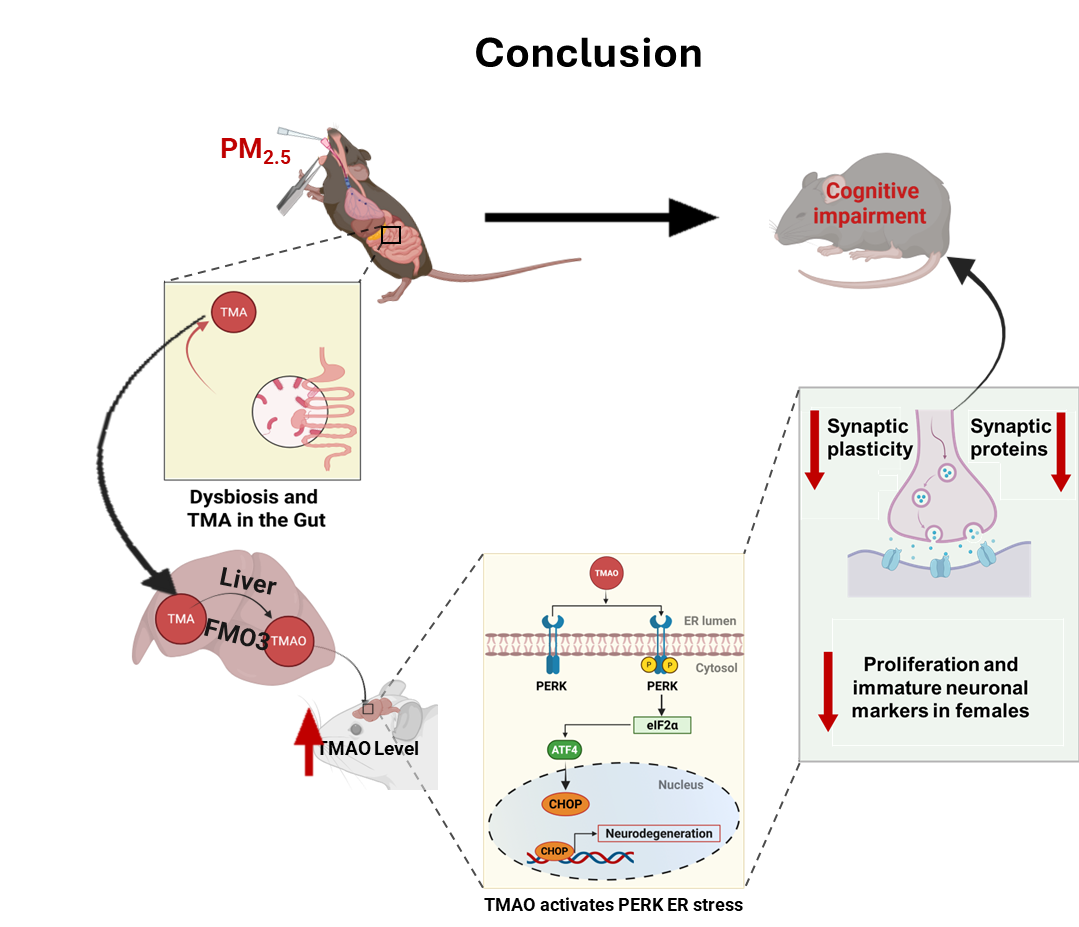
**Graphical Abstract**

**Fig. S8.** Proposed mechanism linking chronic PM2.5 exposure to hippocampal dysfunction and cognitive impairment. Chronic inhalation of PM2.5 perturbs gut microbiota composition, increasing microbial production of trimethylamine (TMA) from dietary precursors. TMA is absorbed and converted in the liver by flavin‑containing monooxygenase 3 (FMO3) to trimethylamine‑N‑oxide (TMAO), leading to elevated systemic and hippocampal TMAO. Increased TMAO in the brain activates endoplasmic reticulum (ER) stress via the PERK → eIF2α → ATF4 → CHOP signaling cascade, promoting neurodegenerative processes. Downstream consequences in the hippocampus include reduced synaptic plasticity and synaptic protein expression and impaired adult neurogenesis (reduced proliferation and immature neuronal markers), with some effects more pronounced in females. Genetic or pharmacological reduction of FMO3/TMAO production (for example, FMO3 disruption or DMB) and hippocampal PERK knockdown mitigate ER stress, preserve synaptic proteins and neurogenic markers, and attenuate PM2.5‑induced cognitive deficits.

**Table S1. List of antibodies used**

| **Antibody** | **Host species** | **Catalog no.** | **Dilution** | **Supplier** |
| --- | --- | --- | --- | --- |
| FMO3 | Rabbit | 17469-1-AP | 1:500 | Proteintech |
| BiP | Rabbit | 3177 | 1:1000 | CST |
| eIF2α | Rabbit | 5324 | 1:1000 | CST |
| ATF-4 | Rabbit | 11815 | 1:1000 | CST |
| PERK | Rabbit | 5683 | 1:1000 | CST |
| Phospho-eIF2α (Ser51) | Rabbit | 9721 | 1:1000 | CST |
| P-PERK (Thr982) | Rabbit | DF7576 | 1:1000 | CST |
| CHOP | Mouse | 2895 | 1:1000 | CST |
| PSD-95 | Rabbit | 2507S | 1:1000 | CST |
| Synaptophysin | Rabbit | 5461S | 1:1000 | CST |
| GluN2A | Rabbit | 4205S | 1:500 | CST |
| GluA1 | Rabbit | 13185s | 1:1000 | CST |
| α-tubulin | Rabbit | 2144s | 1:1000 | CST |
| Ki67 | Rabbit | 9129 | 1:200 | CST |
| Doublecortin | Rabbit | sc-271390 | 1:200 | Santa cruz |
| β-actin | Rabbit | 4967S | 1:1000 | CST |
